# Comparative analysis defines a broader FMRFamide-gated sodium channel family and determinants of neuropeptide sensitivity

**DOI:** 10.1101/2022.03.23.485451

**Authors:** Mowgli Dandamudi, Harald Hausen, Timothy Lynagh

## Abstract

FMRFamide and similar neuropeptides are important physiological modulators in most invertebrates, but the molecular basis of FMRFamide activity at its receptors is unknown. We therefore sought to identify the molecular determinants of FMRFamide potency in one of its native targets, the excitatory FMRFamide-gated sodium channel (FaNaC) from gastropod mollusks. Using molecular phylogenetics and electrophysiological measurement of function, we identified a broad FaNaC family that includes mollusk and annelid channels gated by FMRFamide, FVRIamides, and/or Wamides (or myoinhibitory peptides). A comparative analysis of this broader FaNaC family and other channels from the overarching DEG/ENaC superfamily, incorporating mutagenesis and experimental dissection of function, identified a pocket of amino acid residues that determines activation of FaNaCs by neuropeptides. Although this pocket has diverged in distantly related DEG/ENaC channels that are activated by other ligands, such as mammalian acid-sensing ion channels, we show that it nonetheless contains residues that determine enhancement of those channels by similar peptides. This study thus identifies amino acid residues that determine FMRFamide activity at FaNaCs and illuminates evolution of ligand recognition in one branch of the DEG/ENaC superfamily of ion channels.

## Introduction

Neuropeptides are a large, diverse group of intercellular signaling molecules found in most animals (1). They commonly range in length from four to 40 amino acid residues and are generally the cleavage product of a long pro-peptide containing multiple shorter peptides between peptidase cleavage sites. By diffusing to and binding their receptor targets on other cells, which are most often G-protein coupled receptors (GPCRs), neuropeptides modulate numerous physiological functions. These include neuronal excitability, ciliary beating and locomotion, and muscle contraction, in both cnidarians (e.g. sea anemones and jellyfish) and bilaterians (e.g. worms, flies, and mammals) (2-5).

The 4-mer Phe-Met-Arg-Phe-amide (FMRFa) is a notable example. It is broadly expressed in neurons of invertebrate bilaterians, making it a commonly used marker of the nervous system in immunohistochemical experiments (6-8). Work in selected model bilaterians such as the ecdysozoans *Drosophila melanogaster* and *Caenorhabditis elegans* and the spiralian *Platynereis dumerilii* (an annelid) has begun identifying physiological responses mediated by FMRFa, cognate peptide/receptor pairs, and how these differ across different animals (9-12). Gastropod mollusks (slugs and snails - another group of spiralians) additionally possess an FMRFa-gated sodium channel (FaNaC), a receptor that mediates rapid neuronal and cardiac excitation upon FMRFa binding (13-15).

FaNaCs are members of the degenerin/epithelial sodium channel (DEG/ENaC) superfamily of trimeric sodium channels (14). In contrast to FaNaCs, the DEG/ENaC channels found in mammals are gated by increased proton concentrations (acid-sensing ion channels, ASICs), gated by bile acid (bile acid-gated channels, BASICs), or are constitutively active (ENaCs) (16,17). However, the binding of FMRFa or similar RFamides to ASICs has been shown to enhance proton-gated currents (18), perhaps offering a glimpse of their shared evolutionary history with FaNaCs and also raising interest in peptide-based pharmacological modulators of mammalian channels (15,19). Knowledge of the molecular basis for FMRFa activity at its receptors would therefore offer insight into the role of neuropeptides and receptors in the evolution of bilaterian physiology and also enable the rational design of novel pharmacological modulators of mammalian receptors (20-22). But unfortunately, this knowledge is lacking.

We therefore sought to establish the molecular determinants of FMRFa activity in FaNaCs. Although previously described in only gastropod mollusks, we sought to characterize the FaNaC family in more depth and therefore considered closely related genes from other spiralians, such as annelids and platyhelminths, using phylogenetics, heterologous expression, and electrophysiological experiments. The resulting picture of the FaNaC family allowed us to experimentally probe the molecular basis of FMRFa sensitivity in the broader family using site-directed mutagenesis. Together, our results identify key determinants of channel activation and modulation by FMRFa and other neuropeptides and offer insight into the evolution of the FaNaC family.

## Results

### Phylogenetic and experimental characterization of the FaNaC family

To identify the amino acid residues that determine FMRFa activity at FaNaCs, we considered comparing the amino acid sequences of previously characterized gastropod FaNaCs with other closely related channels of different function. However, gastropod FaNaCs comprise a narrow branch of the overarching DEG/ENaC superfamily, and sequence differences between this small number of channels and other DEG/ENaC channels would be numerous, making the list of candidate amino acid residues long and impractical. We therefore sought a broader view of the FaNaC family and questioned if it extends beyond gastropods. To better estimate the breadth of the FaNaC family within the DEG/ENaC superfamily, we assembled DEG/EnaC sequences from numerous bilaterians and several other lineages and generated a maximum likelihood phylogeny of 544 non-redundant sequences (Fig. 1). Previously characterised FaNaCs from gastropods such as *Helix aspersa* (14) appear in a well-supported clade, hereafter the “FaNaC branch” (Fig. 1A, blue). Notably, this branch also includes closely related sequences from other mollusks and from annelids and brachiopods, tentatively suggesting that FaNaCs occur in numerous spiralian animals. But the only non-gastropod sequences from this branch that have been previously characterized are four from the annelid *Platyenereis dumerilii*, and when heterologously expressed, these did not respond to FMRFa, although one was activated by larger “Wamides”, such as GWKQGASYSWa (23) (“MGIC *Platynereis dumerilii*”, pink in Fig. 1A).

**Figure 1.**
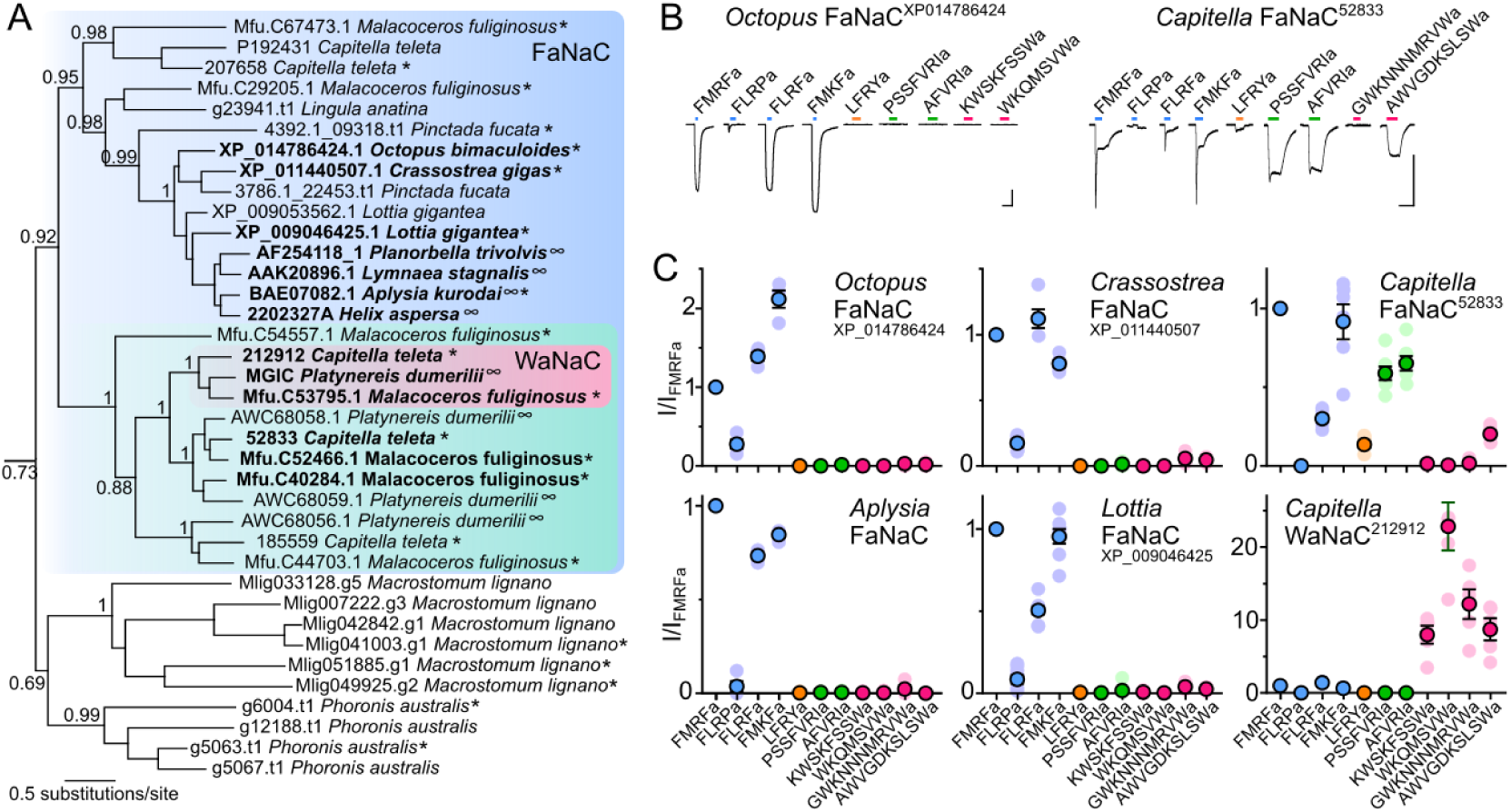
Phylogenetic and experimental characterization of the FaNaC family. (**A**) FaNaC branch from DEG/ENaC phylogeny (full phylogeny in Fig. S1). *Genes tested here. ∞Genes tested elsewhere. Genes in bold encode peptide-gated channels. aLRT support for key branches is shown. (**B**) Example peptide-gated currents at *Xenopus* oocytes expressing indicated genes. All peptide applications 100 μM, except for some at *Capitella* FaNaC: FMRFa, FLRPa, FLRFa, and FMKFa (3 μM); LFRYa (30 μM). Scale bars: x, 10 s; y, 5 μA. **(C)** Mean (± SEM) normalized current response to different peptides (concentrations as in (**B**)) at oocytes expressing indicated channels. Lighter symbols are individual data points, n = 4-7.

We next sought experimental evaluation of the broader FaNaC family seen in our phylogeny. To this end, we injected *Xenopus laevis* oocytes with cRNA of 20 uncharacterized genes from this or closely related branches (asterisks in Fig. 1A) and measured electrophysiological responses to FMRFa and a selection of neuropeptides encoded by the RFamide, FVRIamide, and Wamide pro-peptides from these animals (Fig. 2). To compare these with canonical gastropod FaNaCs, we performed similar experiments on *Aplysia kurodai* FaNaC (24). Genes from the cephalopod *Octopus bimaculoides*, the bivalve *Crassostrea gigas*, and the gastropod *Lottia gigantea* encoded FaNaCs similar to the gastropod *Aplysia* FaNaC, evident in large inward currents in response to both FMRFa and closely related peptides (Fig. 1B,C). FMKFa and FLRFa, which in addition to FMRFa are encoded by the *Octopus* FMRFa pro-peptide (Fig. 2), activated larger currents than FMRFa at *Octopus* FaNaC, and FLRFa also activated slightly larger currents than FMRFa at *Crassostrea* FaNaC (Fig. 1B,C). Interestingly, we observed that two genes from the annelid *Capitella teleta*, 52833 and 212912, also encoded channels gated by RFamides, FVRIamides, and/or Wamides (Fig. 1B,C). This indicates that annelid genes from the FaNaC branch also encode FMRFa-and other peptide-gated channels. One of these, *Capitella* 52833, also showed small currents in response to LFRYa, a product of the RYamide pro-peptide from *Capitella* (Fig. 1B,C). Together, these data describe a broader FaNaC family than previously realized and show that neuropeptides from FMRFa and/or other pro-peptides activate FaNaCs in various mollusks and in annelids.

**Figure 2.**
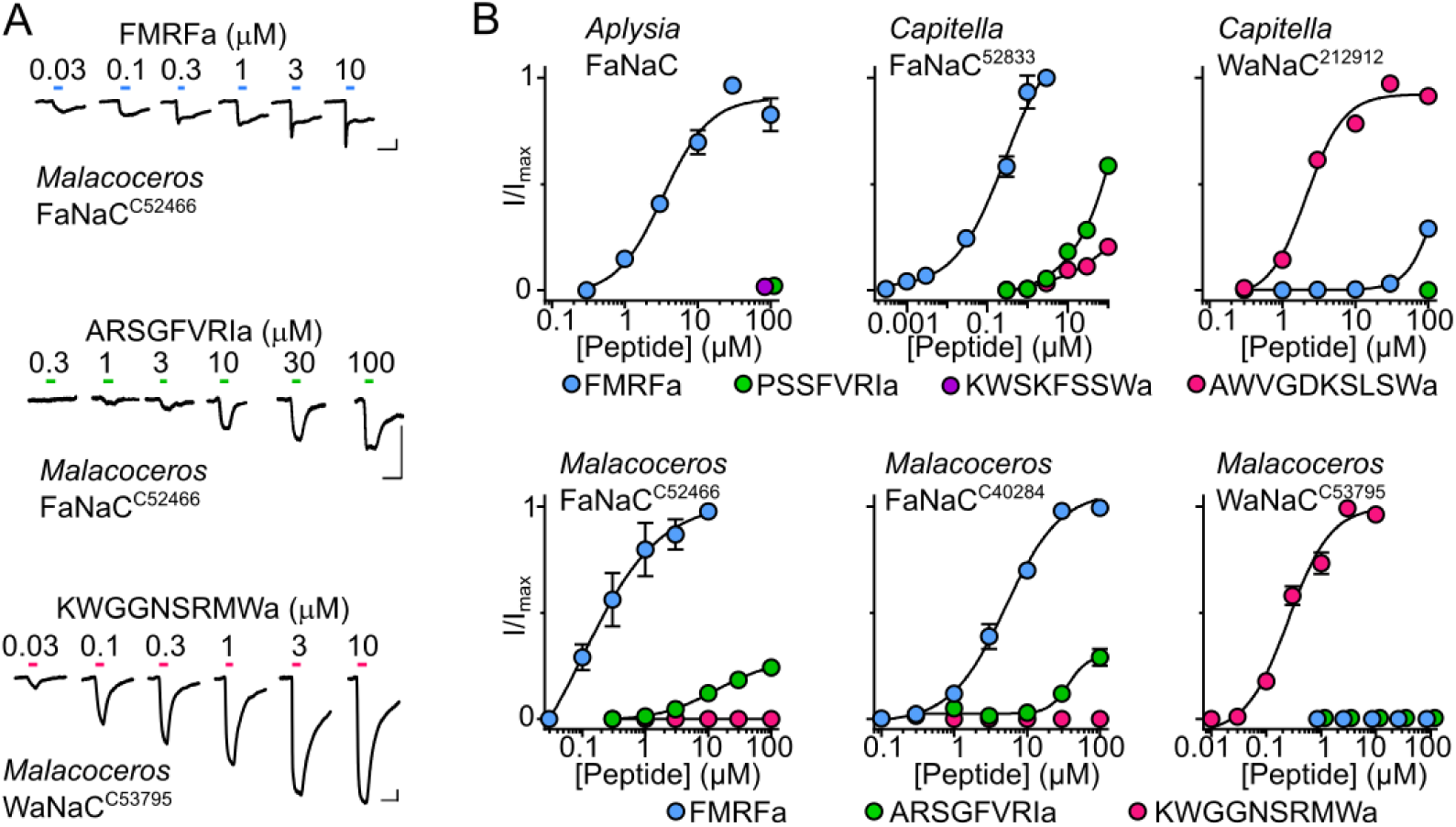
Neuropeptide sensitivity of different members of the FaNaC family. (**A**) Example currents in response to increasing concentrations of FMRFa, ARSGFVRIa, or KWGGNSRMWa at oocytes expressing indicated annelid (*Malacoceros fuliginosus*) channels. Scale bars: x, 10 s; y, 1 μA. (**B**) Mean (± SEM, n = 4-5) normalized current responses to increasing concentrations of neuropeptides (as indicated) at FaNaCs and WaNaCs.

We also tested three platyhelminth (*Macrostomum lignano*) and two phoronid (*Phoronis australis*) genes that appear in a small branch closely related to the FaNaC branch (lower clade in Fig. 1A). We injected oocytes with these RNAs either alone or in combination, in case of obligate heteromerization, but at these oocytes, we observed no response to FMRFa or YMRFa (from *Macrostomum* pro-peptide Mlig001796.g3), FVRIa (from *Phoronis* pro-peptide DN252244_c0_g1_i1), or GRTNNKNVFRWa or AYGSMPWa (both from *Macrostomum* pro-peptide Mlig032062.g1; n = 4-6). Consistent with the phylogenetic relationships we observe, this tentatively suggests that this branch is functionally distinct from the FaNaC branch. Whether this reflects the emergence of FaNaCs after the lineage to Annelida + Brachiopoda + Molluska split from the lineage to Platyhelminthes (and were lost in Phoronida) or whether FaNaCs emerged earlier and FMRFa sensitivity was lost in these platyhelminth and phoronid channels would require experiments on more channels. Furthermore, we cannot rule out that certain channels simply express poorly in our heterologous system, which could also relate to one mollusk and six annelid genes from the FaNaC branch itself, which showed no currents in response to FMRFa and other RFamides or to LFRYa, FVRIamides, or Wamides (n = 3-7; asterisks/non-bold font in Fig. 1A).

Activation of annelid FaNaCs by RFamides, FVRIamides, and Wamides contrasts with activation of mollusk FaNaCs by only RFamides (14,23,24) (Fig. 1C). We investigated this further by testing additional channels from this clade of annelid genes (green in Fig. 1A), also utilizing transcripts from another annelid, *Malacoceros fuliginosus. Capitella* 52833 and *Malacoceros* C52466 and C40284 were more potently activated by FMRFa than cognate Wamides and FVRIamides (Fig. 2A,B), and we therefore refer to these channels as FaNaCs. The half-maximum effective concentration (EC_50_) of FMRFa was 350 ± 150 nM (n = 7) at *Capitella* FaNaC^52833^ and 230 ± 60 nM (n = 5) and 11 ± 3 μM (n = 8) and at *Malacoceros* FaNaC^C52466^ and FaNaC^C40284^, respectively. In contrast, *Capitella* 212912 and *Malacoceros* C53795 were more potently activated by cognate Wamides than by FMRFa and FVRIamides (Fig. 2A,B), much like the previously characterized *Platynereis* MGIC from the same clade (pink in Fig. 1A, (23)), and we therefore refer to these as Wamide-gated sodium channels (WaNaCs). The AWVGDKSLSWa EC_50_ at *Capitella* WaNaC^212912^ was 2 ± 0.2 μM (n = 8) and the KWGGNSRMWa EC_50_ at *Malacoceros* WaNaC^C53795^ was 340 ± 100 nM (n = 9). These results identify within the FaNaC branch a clade of annelid genes that evolved sensitivity to neuropeptides from other pro-peptides.

### Comparison of FaNaCs with other DEG/ENaCs identifies molecular determinants of FMRFa sensitivity

With this broader view of FaNaC sequence and function, we returned to the aim of comparing amino acid sequences to identify amino acid residues that determine activation by FMRFa. An amino acid sequence alignment comparing the FaNaC family with other DEG/ENaC channels, focusing on the putative extracellular and upper transmembrane domains of the protein, where ligand binding is likely to occur, identified 43 amino acid residues that are reasonably conserved in FMRFa-gated channels but divergent in other DEG/ENaCs (Fig. 3A, Fig. 3). These 43 FaNaC-specific residues are spread throughout the extracellular domain, according to a homology-based structural model of *Aplysia kurodai* FaNaC (Fig. 3B). We took *Aplysia* FaNaC as a representative FaNaC, and these 43 residues were individually (or together with one or two vicinal residues, for efficiency) mutated to an equivalent residue from channels not gated by FMRFa, generating 30 *Aplysia* FaNaC mutants (e.g. A319Q in Fig. 3A). FMRFa potency at wild-type (WT) and mutant channels was then measured in electrophysiological experiments. Current responses to 3 μM and 30 μM FMRFa were analysed, giving an “I_3μM_/I_30μM_ ratio” of FMRFa potency, which was 0.64 ± 0.04 for WT (n = 14, Fig. 3C,D), consistent with the FMRFa EC_50_ of 3.4 ± 0.4 μM (n = 5) from Fig. 2B. 15 mutants showed unchanged or slightly increased FMRFa potency (dark blue in Fig. 3D), and eight mutants showed slightly decreased potency (light blue in Fig. 3D). The remaining seven mutants were not significantly activated by 3 or 30 μM FMRFa (Fig. 3C, white in Fig. 3D), indicating substantial decreases in potency and highlighting these seven positions as potential determinants of FMRFa activity at FaNaCs.

**Figure 3.**
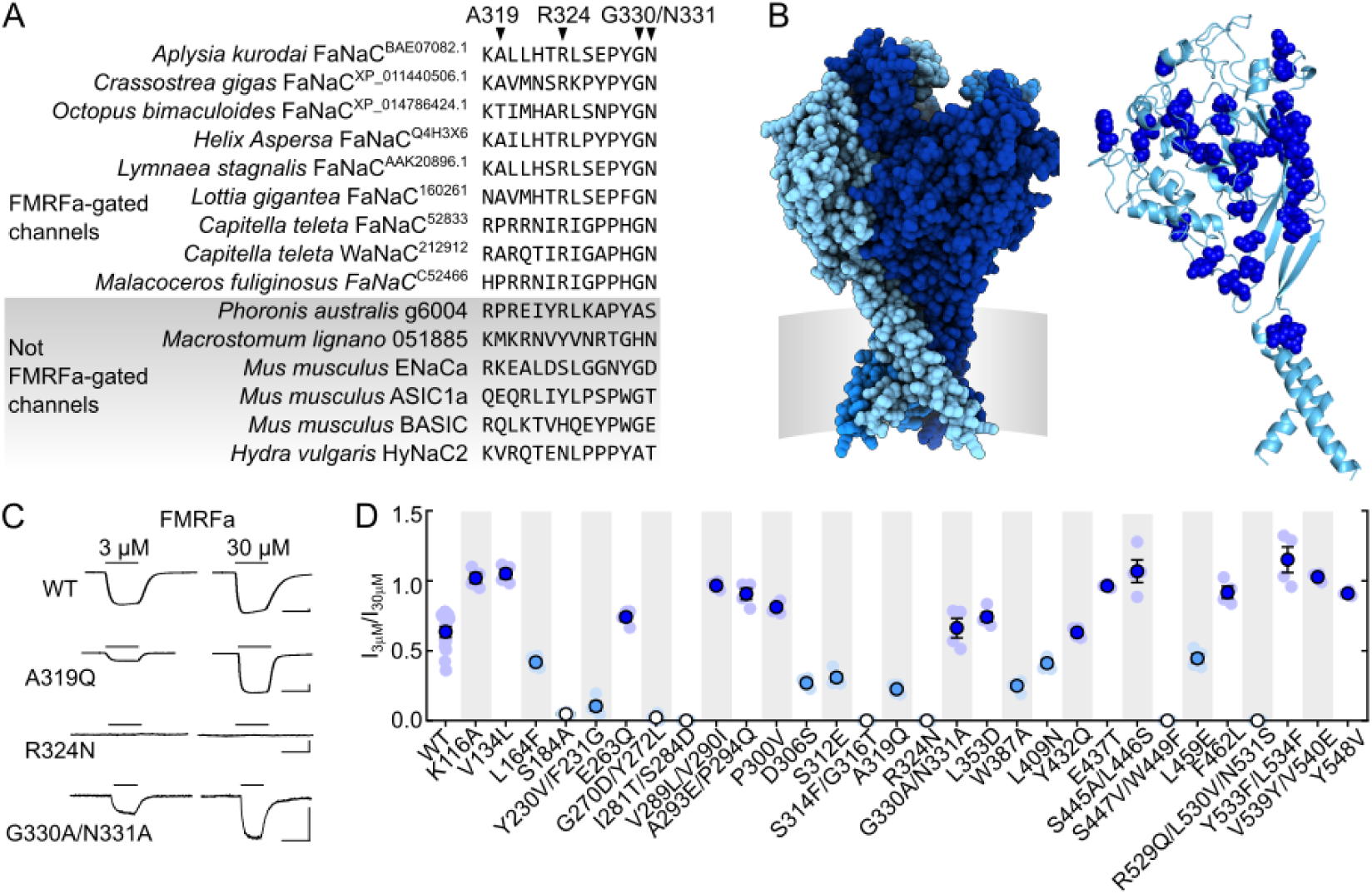
Comparative analysis of amino acid residues determining FMRFa sensitivity. (**A**) Part of amino acid sequence alignment comparing verified FMRFa-gated channels with other channels (complete alignment in Fig. 3). Examples of FaNaC-specific amino acid residues, i.e. those that are conserved – either completely or in terms of physico-chemical properties – in FaNaCs but different in other channels, are indicated by arrowheads. (**B**) *Left, Aplysia kurodai* FaNaC homology model. Different subunits in different shades of blue, approximate position of cell membrane in grey. *Right*, 43 FaNaC-specific amino acid residues (dark blue spheres) in a single subunit of model. (**C**) Example current responses to FMRFa at oocytes expressing WT or mutant *Aplysia kurodai* FaNaC channels. Scale bars: x, 5 s; y, 1 μA. (**D**) Mean (± SEM) FMRFa potency (3 μM FMRFa-gated current amplitude/30 μM FMRFa-gated current amplitude: “I_3μM_/I_30μM_”) was compared by one-way ANOVA with Dunnett’s multiple comparisons test: dark blue, not significantly different to WT or significantly greater than WT; light blue, significantly less than WT (*P* < 0.05); white, not significantly different from zero. Individual data points shown as faint symbols, n = 3-14.

Of the seven mutants that showed a loss of FMRFa potency, two involved single substitutions (S184A and R324N), and the other five involved multiple substitutions (G270D/Y272L, I281T/S284D, S314F/G316T, S447V/W449F, and R529Q/L530V/N531S). We therefore clarified which of the substitutions underlay the functional effect in these five mutants by making additional single-substitution *Aplysia* FaNaC mutants. We did the same for Y230V/F231G, as this double-mutant showed the greatest decrease in FMRFa potency among the “moderate” mutants in Fig. 3D (light blue). In five of these six cases, this clearly identified a single residue whose mutation decreases FMRFa potency (Fig. 4A,B), namely F231G, I281T, G316T, S447V, and R529Q. In contrast, the single G270D and Y272L mutations only moderately decreased FMRFa potency (Fig. 4B), and we did not analyze these further. Full FMRFa concentration-response experiments on the loss-of-function single-mutants revealed greater decreases in FMRFa potency caused by F231G (small currents activated by 100 μM), I281T, G316T, R324N, S447V, and R529Q (negligible, if any, current activated by 100 μM) than by S184A (EC_50_ ≈ 30 uM, Fig. 4C,D).

**Figure 4.**
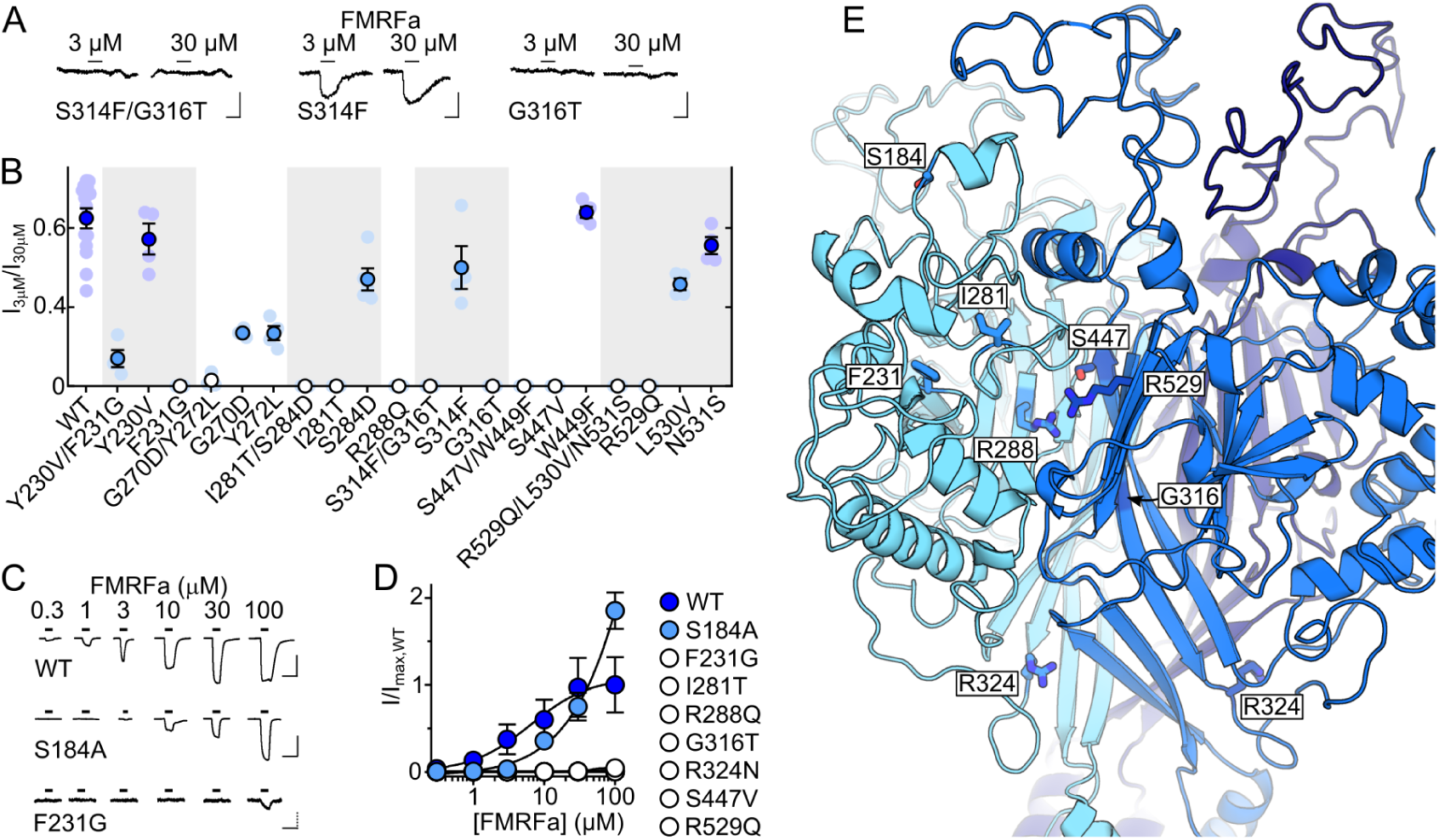
Isolation of individual determinants of FMRFa sensitivity in *Aplysia* FaNaC. (**A**) Example current responses to FMRFa at oocytes expressing mutant *Aplysia kurodai* FaNaC channels. Scale bars: x, 5 s; y, 200 nA. (**B**) Mean (± SEM) FMRFa potency (3 μM FMRFa-gated current amplitude/30 μM FMRFa-gated current amplitude: “I_3μM_/I_30μM_”), WT repeated from Fig. 3. Dark blue, not significantly different to WT; light blue, significantly less than WT (*P* < 0.05); white, not significantly different from zero. Individual data points as faint symbols, n = 4-14. (**C**) Example responses to increasing FMRFa concentrations at indicated *Aplysia* FaNaC mutants. Scale bars: x, 10 s; y, 10 μA (WT and S184A) or 500 nA (F231G). (**D**) Mean (± SEM, n =3-6) normalized current amplitude in response to increasing FMRFa concentrations. Each individual data point was normalized to mean maximum current amplitude at WT (n = 3) on the same day. (**E**) Extracellular domain of *Aplysia* FaNaC homology model showing selected residues (sticks) at the interface of two adjacent subunits (one cyan, one blue).

Seeking a potential explanation for the large effects of these mutations on FMRFa potency, we examined their location in a homology-based structural model of *Aplysia* FaNaC. Remarkably, F231, I281, G316, S447, and R529 all line a pocket at the interface of adjacent subunits in the extracellular domain of the channel (Fig. 4E). As the side chain of an additional residue, R288, auspiciously orients directly into this pocket, we also generated an R288Q mutant, and observed that this mutation also abolished FMRFa potency (Fig. 4B,D). We thus arrived at a list of seven major determinants of FMRFa activity in *Aplysia* FaNaC: F231, I281, R288, G316, S447, and R529, which line the extracellular pocket at the interface of adjacent subunits, and R324, which is in a loop immediately downstream of the β-strand containing G316 (Fig. 4E).

### Molecular determinants of neuropeptide activity in mollusk and annelid FaNaCs and WaNaCs

We next tested if the close phylogenetic relationship of mollusk and annelid FaNaCs and WaNaCs is reflected in similar molecular determinants of neuropeptide activity at these channels. We mutated *Capitella* FaNaC^52833^ and *Capitella* WaNaC^212912^ at positions equivalent to the seven crucial residues identified in *Aplysia* FaNaC above and tested mutant channels for responses to FMRFa or AWVGDKSLSWa (Fig. 5A). In *Capitella* FaNaC^52833^, only R278Q and G306T mutations, equivalent to *Aplysia* FaNaC R288 and G316, abolished FMRFa-gated currents (green in Fig. 5B). In *Capitella* WaNaC^212912^ R255Q, R291N, and R472Q mutations, equivalent to *Aplysia* FaNaC R288, R324, and R529, abolished AWVGDKSLSWa-gated currents (pink in Fig. 5B). Thus, the glycine residue in β-strand 9 (G316 in *Aplysia* FaNaC) seems crucial for FMRFa activation of FaNaCs, the arginine residue in β-strand 7 (R288 in *Aplysia* FaNaC) is crucial for neuropeptide activation of both FaNaCs and WaNaCs, and generally, arginine residues make a substantial contribution to channel sensitivity to neuropeptides.

**Figure 5.**
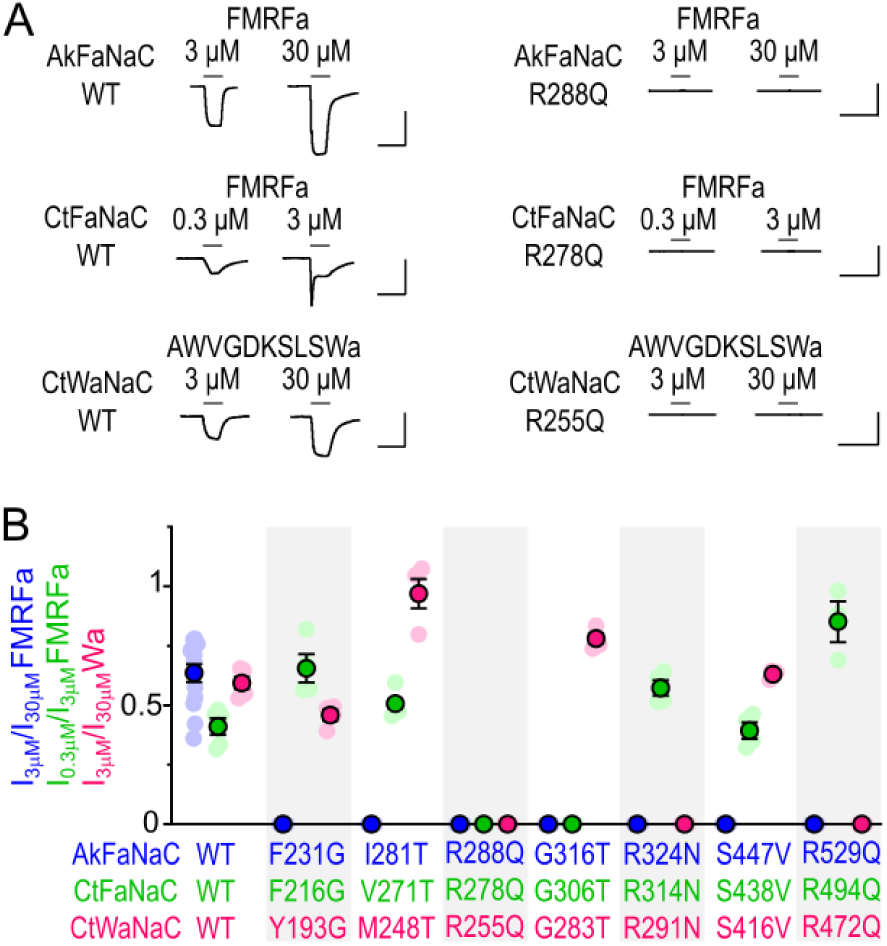
Determinants of neuropeptide activity in different FaNaCs and WaNaCs. (**A**) Example current responses to indicated neuropeptide at oocytes expressing WT or mutant *Aplysia* FaNaC (AkFaNaC), *Capitella* FaNaC^52833^ (CtFaNaC), or *Capitella* WaNaC^212912^ (CtWaNaC). Scale bars: x, 10 s; y, 5 μA. (**B**) Average (± SEM, n = 4-5) FMRFa potency (I_3μM_/I_30μM_ at *Aplysia* FaNaC, I_0.3μM_/I_3μM_ at *Capitella* FaNaC) and AWVGDKSLSWa potency (I_3μM_/I_30μM_ at *Capitella* WaNaC), grouped according to the equivalent amino acid residue.

### Conserved determinants of neuropeptide sensitivity in distantly related DEG/ENaC channels

Acid-sensing ion channels (ASICs) are members of the DEG/ENaC superfamily that are distantly related to FaNaCs (Fig. S1). ASICs are found in numerous bilaterians, including annelids, but are best described in mammals (25). Mammalian ASICs are closed at pH ≥ 7.4 and transiently activated by e.g. pH 5, but if resting channels are exposed to very small drops in pH, e.g. to pH 7, they enter a desensitized state, and subsequently pH 5 activates little, if any, current (Fig. 4B). However, in the presence of the synthetic FMRFa analogue FRRFa, pH 7 induces less desensitization, and subsequently pH 5 activates substantial current (19) (Fig. 4C). FRRFa thus enhances ASICs, and finally, we questioned if this enhancement is determined by amino acid residues in a similar pocket to that identified in FaNaCs and WaNaCs above.

Although we used mouse sequences in our earlier amino acid sequence analyses, we employed rat ASIC1a in our experiments, because we had the clone in our laboratory, and rat and mouse ASIC1a differ at only one amino acid position in the intracellular domain. We measured FRRFa enhancement of pH-gated currents in rat ASIC1a WT and mutants carrying alanine substitutions at six of the seven positions addressed above: T239A (equivalent to *Aplysia* FaNaC I281), K246A (FaNaC R288), S274A (FaNaC G316), Y282A (FaNaC R324), V377A (FaNaC S447), and V405A (FaNaC R529; full alignment in Fig. 3). We did not pursue ASIC1a G192 (FaNaC F231), as it was precisely the loss of a side chain in the *Aplysia* FaNaC F231G mutant that abolished FMRFa activity. At WT and four of the six mutant ASICs, FRRFa showed 4-to 12-fold enhancement of pH-gated currents (WT, 10 ± 3, n = 7; Fig. 6A,B). Although not significantly less than WT (one-way ANOVA with Dunnett’s multiple comparisons test), S274A mutant channels were enhanced only 2 ± 0.3-fold (n = 6), and K246A channels were not ostensibly enhanced at all (n = 6, Fig. 6A,B). Thus, K246 in rat ASIC1a is crucial for FRRFa activity, much like the equivalent R288/R278/R255 residue in FaNaCs and WaNaCs is crucial for FMRFa/Wamide activity.

**Figure 6.**
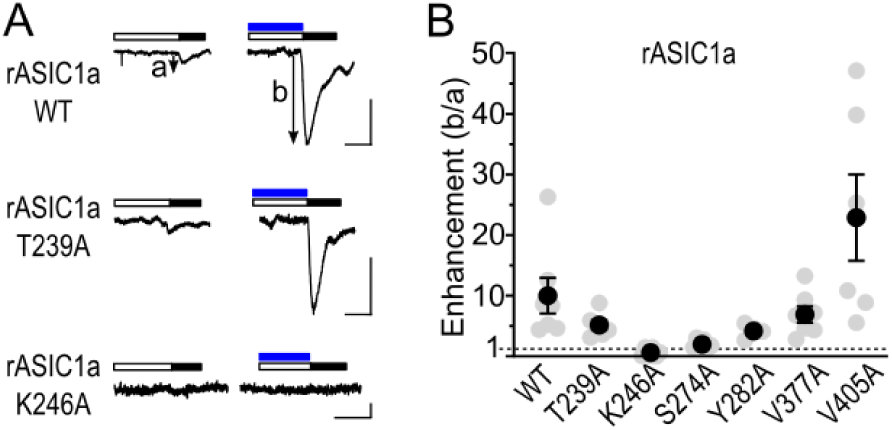
Determinants of FRRFa modulation of rat ASIC1a. (**A**) pH 5.3 (filled black bars) -gated currents after pre-incubation in desensitizing pH (unfilled bars) alone or in 50 μM FRRFa (blue bars) at oocytes expressing WT or mutant rat ASIC1a (rASIC1a). Scale bars: x, 10 s; y, 200 nA. Desensitizing pH described in Fig. 4. (**B**) Mean (± SEM, n = 4-7) enhancement (current_b_/current_a_ from (**A**)) of desensitized current amplitude by 50 μM FRRFa at indicated rat ASIC1a mutants.

## Discussion

We sought to establish the molecular determinants of FMRFa activity at FaNaCs by a comparative analysis of the FaNaC family. Our results lead to two major conclusions. Firstly, FaNaCs are found in multiple spiralian lineages, and in certain lineages, they have evolved sensitivity to other peptides. Secondly, a pocket in the extracellular domain of FaNaCs and other DEG/ENaC channels contains several amino acid residues whose mutation abolishes FMRFa and/or other neuropeptide activity.

### FaNaC evolution

Our combined phylogenetic and experimental analysis offers good evidence for a broader FaNaC family including mollusk, brachiopod, and annelid genes (blue in Fig. 1A). Within the FaNaC family, we see a well-supported clade of annelid genes encoding channels gated by FMRFa and/or other peptides from RFamide, FVRIamide, RYamide, and Wamide pro-peptides (green in Fig. 1A), and a well-supported clade of mollusk and brachiopod genes encoding channels gated by FMRFa and similar peptides from only the RFamide pro-peptide (upper clade in Fig. 1A). This suggests that the last common ancestor of annelids and mollusks had at least one gene encoding a FaNaC. At least two rounds of FaNaC gene duplications are evident in annelids, with one of the copies gaining sensitivity also to other neuropeptides such as Wamides, FVRIamides, and LFRYamide. Some of these genes, such as *Malacoceros* WaNaC^C53795^, have lost sensitivity to FMRFa.

More precise conclusions, especially on the early diversification of FaNaCs in annelids and mollusks, would need more extensive phylogenetic sampling and functional tests of the annelid and brachiopod genes (toward the top of Fig. 1A) that we were unable to characterize and that did not fall into well supported clades. The fact that the closest relatives of the FaNaC family in our tree, genes from platyhelminths and phoronids, do not appear to form FMRF-gated channels tentatively suggests that FaNaCs as we know them emerged in an early ancestor of annelids + brachiopods + mollusks, after this lineage split from the lineage to platyhelminths. However, this interpretation suffers from (a) poor branch support and (b) the possibility that genes we have not tested – in the platyhelminth/phoronid clade and in various other clades – could form FMRFa-gated channels. The fact that such diverse DEG/ENaCs as FaNaCs, ASICs and HyNaCs are sensitive to RFamides, whether activated or only modulated, also suggests that peptide-sensitive channels may be “hiding” in other branches of the DEG/ENaC tree.

### Molecular determinants of FMRFa sensitivity in FaNaCs and other DEG/ENaC channels

Our results identify a conserved arginine residue in FaNaCs and WaNaCs that is required for channel activation by FMRFa and Wamides. Six other conserved FaNaC/WaNaC residues also contribute to FMRFa or Wamide potency, but only in certain channels. According to our homology-based structural model of *Aplysia* FaNaC, six of the seven determinants of potency line a pocket formed by the interface of adjacent subunits in the channel’s extracellular domain. The crucial arginine side chain (R288 in *Aplysia* FaNaC) orients from the left subunit into the middle of the pocket.

Knowledge of the structure of DEG/ENaC channels derives mostly from X-ray crystallography of chick ASIC1 (e.g. (26)), on which our *Aplysia* FaNaC model is based. Our model suffers from low sequence identity with the template (21%) and long loops between extracellular β-strands relative to the template. Therefore, the location and orientation of F231 and I281 in our model is questionable. However, the location of R288, G316, S447, and R529 in β-strands forming the interfacial pocket, and even R324 in a “thumb domain” loop downstream of G316 (26), seems more reliable: throughout the DEG/ENaC family these segments are more conserved, have less indels, and are also close to conserved cysteine residues that stabilize structure via disulfide bonds (Fig. 3). The crucial arginine, in particular, may have a similar orientation and functional role in neuropeptide sensitivity in diverse DEG/ENaCs. It was not identified in our initial comparative analysis, because rather than being FaNaC-specific, a basic arginine or lysine residue occurs at this position in numerous DEG/ENaCs (Fig. 3). Our observation that FRRFa enhancement of rat ASIC1a was abolished by the K246A mutation is consistent with a conserved role of this basic residue in neuropeptide sensitivity.

Although few previous studies have probed the determinants of RFamide sensitivity in DEG/ENaCs, we note certain reports of mutations that affect RFamide activity at gastropod FaNaCs and mammalian ASICs. The substitution of a 32-amino acid segment (including the residue equivalent to *Aplysia* FaNaC F231) from *Helix aspersa* FaNaC (FMRFa EC_50_ = 3 μM) into *Helisoma trivolvis* FaNaC (FMRFa EC_50_ = 100 μM) conferred 3 uM FMRFa sensitivity on the latter (27)). Rat ASIC1a S274 (equivalent to *Aplysia* FaNaC G316), whose mutation to alanine decreased FRRFa enhancement of ASIC1a in our hands, was previously implicated in the different levels of FMRFa enhancement of rodent compared to human ASIC1a (19) and is reasonably close to several other residues whose mutation in human ASIC1a decreases FRRFa enhancement (28).

### Potential roles of extracellular pocket in neuropeptide activity

Residues in the extracellular pocket of FaNaC could determine peptide activity by forming the binding site for peptides or by mediating conformational changes required for channel opening in response to peptide binding elsewhere. Our experiments do not distinguish between these possibilities, but the interfacial, extracellular location of the pocket is reminiscent of binding sites in other ligand-gated ion channel families. In trimeric ATP-gated channels, phylogenetically distinct from DEG/ENaCs but structurally similar and with a similarly sized agonist, ATP binds in the extracellular domain at the interface of adjacent subunits, causing conformational changes that lead to channel opening (29). Similarly, in pentameric ligand-gated ion channels, such as excitatory nicotinic acetylcholine receptors, agonist binding to the interface of adjacent subunits sees the left subunit close around the agonist pulling the lower parts of the receptor into an open-channel state (30). (The remaining ligand-gated ion channel superfamily of tetrameric ionotropic glutamate receptors is distinct, in that their structure consists of an extracellular agonist binding domain contained within each subunit, derived from a soluble substrate binding protein in bacteria (31).)

We speculate that RFamides and Wamides bind to the interfacial pocket, where the two aromatics of the neuropeptide could form hydrophobic interactions with F231 or I281 or cation-π interactions with R288 or R529. Similarly, the basic arginine or lysine side chain in most of these neuropeptides could engage with hydrophilic moieties in the channel pocket, such as the S447 side chain or main chain carbonyls from the loops at the top of the pocket, or form cation-π interactions either with the receptor or within the neuropeptide itself. Such interactions are exemplified by the binding mode at other channels of certain peptide toxins and synthetic peptides, some of which, like FMRFa, contain arginine and aromatic side chains in close proximity (32-34). Foreseeably, conformational changes in this part of the extracellular domain of FaNaCs could be mechanically coupled to channel gating via the β-strand containing G316 and, slightly downstream, the loop containing R324, a loop previously implicated in channel gating of diverse DEG/ENaCs (35,36).

### Outlook

As the molecular basis for FMRFa activity at its receptors was previously unknown, we employed a combination of phylogenetics and experimental dissection to investigate FMRFa activity at FaNaCs. In addition to identifying the molecular determinants of FMRFa activity at both gastropod and annelid FaNaCs and WaNaCs, this approach also disentangled evolutionary relationships between these FaNaC family members. Furthermore, it uncovered a molecular and functional link between FaNaCs and more distantly related ASICs. These results enable better prediction of function from sequence and offer a springboard for future studies addressing DEG/ENaC evolution and neuropeptide binding sites in detail.

## Experimental procedures

### Sequence assembly and phylogenetic analysis

To investigate the presence of FMRFa sensitive channels throughout the metazoans, DEG/ENaC amino acid sequences were sought from four mollusks (*Crassostrea gigas, Octopus bimaculoides, Pinctada fucata, Lottia gigantea*), four cnidarians (*Aurelia aurita, Hydra vulgaris, Nematostella vectensis, Stylophora pistillata*), four annelids (*Capitella teleta, Platynereis dumerili, Malacoceros fuliginosus, Helobdella robusta*), one phoronid (*Phoronis australis*), one brachiopod (*Lingula anatina*), one nemertean (*Notospermus geniculatus*), one rotifer (*Branchionus plicatilis*), one platyhelminth (*Macrostomum lignano*), one arthropod (*Araneus ventricosus*), one priapulid (*Priapulus caudatus*), two chordates (*Branchiostoma belcheri, Homo sapiens*), two echinoderms (*Acanthaster planci, Strongylocentrotus purpuratus*), two poriferans (*Amphimedon queenslandica, Oscarella carmela*), one ctenophore (*Mnemiopsis leidyi*), one hemichordate (*Ptychodera flava*). Sequences were obtained through BlastP using the *Aplysia kurodai* FaNaC amino acid sequence (NCBI BAE07082.1) in JGI Capca1 genome (*Capitella teleta*, https://mycocosm.jgi.doe.gov/pages/blast-query.jsf?db=Capca1 (37)), OIST Marine Genomics Unit (*P. fucata, P. australis, N. geniculatus, A. aurita*, https://marinegenomics.oist.jp/gallery), *Macrostomum lignano* genome initiative (*M. lignano*, http://www.macgenome.org/blast/index.html), Compagen Japan (*O. carmela*, http://203.181.243.155/blast.html), our own *Malacoceros fuliginosus* transcriptome resources (38), or NCBI (for all others, https://blast.ncbi.nlm.nih.gov). After an initial alignment using MAFFT v 7.450 in Geneious Prime (Geneious), removal of >90% identical, redundant sequences and removal of obviously incomplete sequences, sequences were re-aligned with MAFFT and poor-aligning segments and sequences larger than 1000 amino acids or shorter than 300 amino acids were removed. The result was an alignment of 544 sequences with 9,829 amino acid positions. This alignment was used to generate a maximum likelihood tree with PhyML (http://www.atgc-montpellier.fr/phyml/ (39)) using a VT+G substitution model chosen by Smart Model Selection in PhyML (40). Branch support was estimated through aLRT SH-like based method (39).

### Molecular biology, chemicals, and peptides

Sequences chosen for cDNA synthesis, cRNA transcription and expression were *C. gigas* XM_011442205.2, *O. bimaculoides* XM_014930938.1, *L. gigantea* XM_009055314.1_160261, *P. fucata* pfu_aug2.0_4392.1_09318.t1, *C. teleta* Capca1_207658, Capca1_52833 (alternative splice variant, Capca1_191096, using fgenesh ab initio models was used), Capca1_185559, and Capca1_212912, *M. fuliginosus* C67473.1, C29205.1, C54557.1, C53795.1, C52466.1, C40284.1, and C44703.1, *M. lignano* Mlig049925.g2, Mlig051885.g1, and Mlig041003.g1, and *P. australis* g6004.t1 and g5063.t1. These were selected based on their position in the FaNaC or adjacent clade (Fig. 1A) and to a lesser extent, which animal lineage they represent. *A. kurodai* BAE07082.1 (*Aplysia* FaNaC) cDNA in a modified pSP64 vector was provided by Prof. Yasuo Furukawa, Hiroshima University, Japan. *Rattus norvegicus* NP_077068.1 (rat ASIC1a) in the pRSSP6009 vector was provided by Prof. Stefan Gründer, RWTH Aachen University. cDNA for the other sequences was commercially synthesized and sub-cloned (Genscript) between SalI and BamHI sites in a pSP64 (polyA) vector (Promega) modified based on (41) and containing a C-terminal cMyc tag (Suplementary Information Text).

Site directed mutagenesis was performed with custom designed primers (Merck) and PCR with Phusion High-Fidelity DNA polymerase (ThermoFisher Scientific) following the supplier’s PCR protocol and using primers designed according to (42). All sequences – mutant and WT – were confirmed by Sanger sequencing of the full insert (Genewiz). cDNAs were linearized with EcoRI, Pdil or XbaI (ThermoFisher Scientific), and cRNA was transcribed with the mMESSAGE mMACHINE SP6 Transcription Kit (ThermoFisher Scientific).

Peptides were custom synthesized by Genscript, with mass was confirmed by electrospray ionization mass spectrometry, ≥95% purity confirmed by reversed-phase high performance liquid chromatography, and trifluoroacetic acid replaced with acetic acid. Salts and other standard chemicals were purchased from Merck.

### Heterologous expression and electrophysiology

Defolliculated stage V/VI oocytes from *Xenopus laevis* were commercially acquired (Ecocyte Biosciences) and stored at 18 °C in 50% (in water) Leibovitz’s L-15 medium (Gibco) supplemented with additional 0.25 mg/ml gentamicin, 1 mM L-glutamine, and 15 mM HEPES, pH 7.6. Oocytes were injected with 2 ng cRNA (wild type or mutant *Aplysia* FaNaC) and 2-60 ng cRNA (wild type *C. gigas* XM_011442205.2, *O. bimaculoides* XM_014930938.1, *L. gigantea* XM_009055314.1_160261, *P. fucata* pfu_aug2.0_4392.1_09318.t1, *C. teleta* Capca1_207658, Capca1_52833, Capca1_185559, and Capca1_212912, *M. lignano* Mlig049925.g2, Mlig051885.g1, and Mlig041003.g1, *P. australis* g6004.t1 and g5063.t1 and *R. norvegicus* NP_077068.1 (rat ASIC1a) and respective mutants). Two-electrode voltage clamp experiments were performed one to three days after injection. The oocyte was placed in an RC-3Z bath (Warner Instruments), and bath solution (NaCl: 96mM, KCl: 2mM, CaCl_2_: 1.8mM, MgCl_2_: 1mM, HEPES: 5mM; pH: 7.5) was perfused continuously and rapidly switched to bath solution containing peptide ligands via the VCS-8-pinch valve control perfusion system (Warner instruments). In initially determining peptide sensitivity of various channels, all peptides were applied at 100 μM, except at *Capitella* FaNaC^52833^ which showed slow washout of FMRFa-gated current and was therefore exposed to 3 μM FMRFa, FLRPa, FLRFa, and FMKFa, 30 μM LFRYa, and 100 μM PSSFVRIa, GWKNNNMRVWa and AWVGDKSLSWa. Peptides were applied at varying concentrations between 30 nM and 100 μM for concentration-response experiments. The oocytes were clamped at -40 mV for rat ASIC1a experiments and -60 mV for all others with an Oocyte Clamp OC-725C amplifier (Warner Instruments), LIH 8+8 digitiser (HEKA Elektronik) at 0.5 or 1 KHz sampling and 100 Hz filtering. Current recordings were analyzed in Clampfit 11.1 (Molecular Devices).

After retrieving current amplitude from pClamp, all data analyses were performed in Prism v9 (GraphPad Software). For peptide or proton concentration-response data, peak current amplitude was plotted against peptide concentrations or pH and fitted with the Hill equation for each recording. These were averaged to give the reported means ± SEM in the main text. For display in figures, a single fit to the average normalized responses (± SEM) is shown. Multiple comparisons were made with one-way analysis of variance with Dunnett’s comparison to a control value (e.g. comparing with WT) or with Tukey’s test for multiple comparisons.

### FaNaC structural model

A rudimentary structural model of *Aplysia kurodai* FaNaC was generated by uploading its protein sequence to the Swiss model server (43), which suggested chick ASIC1a structure PDB 4NYK (44) as an appropriate template and generated a pdb file which is vizualised in Fig. 3 and Fig. 4 using PyMOL software (Schrödinger).

## Data availability

DNA sequences are included in supporting information. Amino acid sequence alignment and maximum likelihood tree are available at lynaghlab.com/resources. Electrophysiological data points are shown where possible and are available from the corresponding author on request.

## Supporting information

This article contains supporting information, including references 38 and 41.

## Acknowledgements

We are grateful to Prof. Yasuo Furukawa (Hiroshima University, Japan) for the *Aplysia kurodai* FaNaC clone, Prof. Stefan Gründer (RWTH Aachen University, Germany) for the rat ASIC1a clone, and to Dr. Josep Martí-Solans (University of Bergen, Norway) and Dr. Valeria Kalienkova (University of Groningen, Netherlands) for helpful discussions on the manuscript.

## Conflict of interest

The authors declare that they have no conflicts of interest with the contents of this article.

## Notes

### Competing Interest Statement

The authors have declared no competing interest.

