## Supporting information for "Comparative analysis defines a broader FMRFamide-gated sodium channel family and determinants of neuropeptide sensitivity"

### Contents

**Supporting figures** ........ **S2**

Figure S1 ........ S2

Figure S2 ........ S4

Figure S3 ........ S5

Figure S4 ........ S6

**Supporting text** ........ **S7**

Oocyte expression vector and novel cDNA inserts ........ S7

Other cDNA constructs ........ S10

**Supporting references** ........ **S11**

### Supporting figures

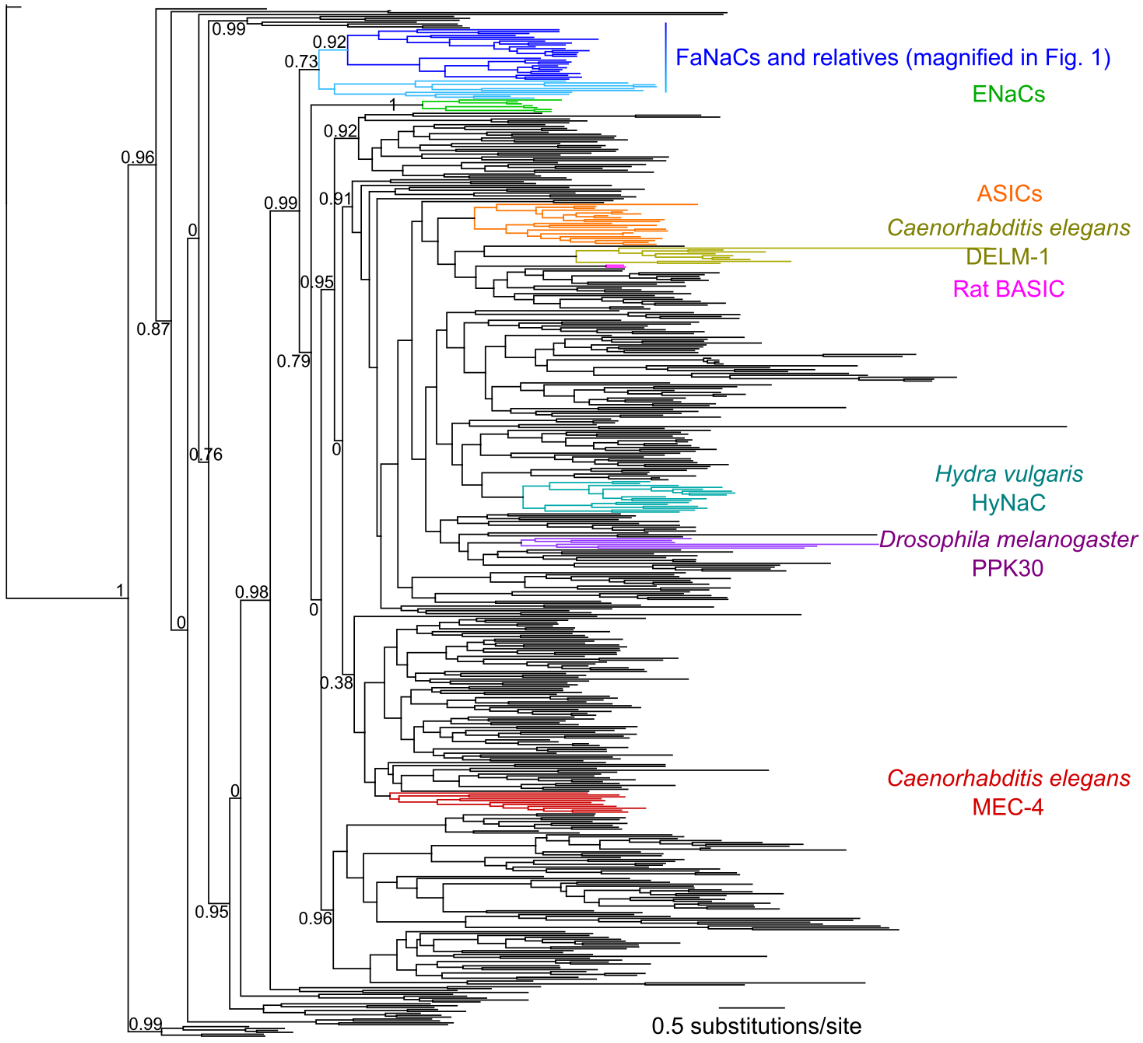


**Figure S1. DEG/ENaC phylogeny.** Unrooted PhyML maximum likelihood tree (ATGC Montpellier Bioinformatics Platform, VT + G calculated as appropriate substitution model) of 544 DEG/ENaC genes from various metazoans. For clarity, only selected branch support values (aLRT SH-like) are shown. Well-known DEG/ENaC sub-families are colored and labeled. Phylogeny is based on MAFFT alignment, both available at https://www.lynaghlab.com/resources.

*Aplysia californica* P08021.3

MRPWCQLALLACLSLKWLTSHVTAESFLCDDSELCENGYLRFGRSMSVEEPHFRLERRSYPPVVYHKRFLRFGRSQEPDIEDYARAIALIESEEPLYRKR
RSADADGQSEKVLHRARREAESEHKSLEEVSPDTKQDVEKRDADDVLDAEKRFMRFGKRFMRFGRGSSDDDESGDDDVQDLTDIGDGLGGEGEVNKRFMR
FGKRFMRFGKREDGEPDKRFMRFGKSMADNDLDKRFMRFGKRFMRFGKSLPDSEVDKRFMRFGKSVDGDVDKRFMRFGKSVDGDVDKRFMRFGKSVDGDV
DKRFMRFGKSVDGDVDKRFMRFGKSVDGDVDKRFMRFGKSVDDDVDKRFMRFGKSVDGDVDKRFMRFGKSVDDAVDKRFMRFGKSVDSDLDKRFMRFGKS
VGSDEVDKRFMRFGKSVGSDEVDKRFMRFGKSLGTDDVNKRFMRFGKSLGTDDVNKRFMRFGKSLGTEDVDKRFMRFGKSLGTDDVDKRFMRFGKSLGTD
DVDKRFMRFGKSLGTEDVDKRFMRFGKSLGTDDVDKRFMRFGKSLGTEDVDKRFMRFGKRFMRFGRSVGDSKYRSASSESVMTTDSKQTTEQATNKS

*Crassostrea gigas* XP_011443786.2

MGTWTYLCLLVAFLLNWFTIETSANDLIDDCYRNPELCQEVGILFGQQQPVDKRFLRFGKRALSGDHYIRFGRNSDDKRFLRFGKRGEQGSVEDDLREAL
NKVIQFKQETGLPLRKRRSADPPLVKDVPEDKDSNSTEKEDSASEKHKRETDEVSEENKRFMRFGRTPAEDDPTYMKRFMRFGRNPDLEKKFMRFGKDGN
EKRFMRFGKREDNDNMVTDDKRFMRFGRDPKDDTLMERFVRNGRSGDDKRFMRFGKRFMRFGKRFDNDGEYDDEDEMGAEKRFMRFGKSGDEEKRFMRFG
KSGDEEKRFMRFGKSGDDEKRFMRFGKSGDEEKRFMRFGKSGDEEKRFMRFGKSGDEEKRFMRFGKSGDEEKRFMRFGKSGDEEKRFMRFGKSGDEEKRF
MRFGKSGDDAKRFMRFGRDPTDKRFMRFGKSV

*Octopus bimaculoides* XP_014775928.1

MSGLSPFSLLIFIFIYCLTVYAATAMSLAQACMESPSMCESISLSHLSSEEGLKSKRFLRPGRALSGDAFLRFGKNGLNLPFEDKRFLRFGRTDRQFEDM
LKEVLQRAENVDNRQKRSTDNIPQSSVQDDSSKITKRNADASDNGMDKRFMKFGKSGDLPYVNEWSDKRFMRFGREPDKRFMRFGKSDDKRFMRFGRNPN
ELEDRLEEGKRFMRFGRGNEEEEKRFMRFGRDPDSKLMGYGNSEEEKRFMRFGRMSDEADAQKRFMRFGRSVDDDKANRLKKSNDQLRTIRMGRSVDDKK
VNSANGDAYLRIGQSDE

*Capitella teleta* 188764

MRIAWMWMVFLLLAVLCLVTEGILLDELCRDADEAKKTSITQFCTTLQKYMDNGNQDAILVLPLSKNQRNSLNDVDDLGAMQNVKRALSDMYFGIRQRRS
GEATNRVRRGGNYIRFGRSAPSSPNPHLADYLMGFGQPDKKAGAYVRFGRSTDAVEKEAEELSESKNEEKRFMRFGKRADDDQKEKKFMRFGRGFMRFGR
SDGLASEDDDKEKRFMRFGKRGDEDEENDEEKRFMRFGKSEGLSEDEAEKEKRFMRFGKREDGEEGGDANEMSEEEKRFMRFGRGFMRFGRDSALEANGK
RYMRFGKKSDTDDEAAASPDKKFMRFGKKFMRFGKRNGEEEDDGKRSGAHAFNPYDQIKRKYMRFGKRADEIPEEKRFMRFGKKFMHFGRDPLADDDESK
DEEEKRFMRFGKKDAANNEVEDTEEETGDTEKRFMRFGKKSSEDAEESPEKKFMRFGKKDSGDQMEDKRFMRFGRSAGAS

*Malacoceros fuliginosus* Mfu.C53703.0.T1

MSVGGVWAAMLLVSMLGSAMGNLATLCARASGPRLQEMCDELILYINEVEGSGPLERSSRMVRDTSSVETLDDGDVTLRDLLESMGQNRRDLLYSIKHRN
KRAGGNYIRFGRSVPDYSSSFGKRDTGAHYIRFGRSTDHVQEQNAMEDAAEKNELVSKVAEEETKAEEEDKRGLRFGKRGDVDDLTEEEKRFMRFGRSTD
DFEGPKRFMRFGKRSDYDDAKRFMRFGKRSDYDDAKRFMRFGKRSDYDDAKRFMRFGKRSDEIDDEDLEEEKRFMRFGKREDEDMDEEKRFMRFGKREND
EDLEEVDADDIEEDKRFMRFGKRESDNDVLGEDKKFMRFGKRFMRFGRDPLDGAKRFMRFGRNNNSGGKRFMRFGKRFMRFGKRSDEDTEDEDEKRFMRF
GKRAEEDEKRFMRFGKKSDEDYGDKRFMRFGKKDGSDEDSLSPKRFMRFGKKDTSDSEEKRFMRFGKKDTTDIDSQTEEDKRFMRFGKRSVSS

Capitella teleta 204015

MWTILLSMVVFVGVVSCSEENQINKRDVSDVAEGDNTFDREARAAIFRYGKRVPIFRYGKRIFRYGKRADQEVEDSDMDLEKRLFRYGRSGSMPTTYKRL
FRYGKRAEDDVEEDDEIEAVKRLFRYGKRGEINTRTAAQQPSVPFRFGERE

*Aplysia californica* XP_005099491.1

MAAVRPTLSYRAMAIPLLLCVVCFMSFCPSAILAEDDASTVKATSPADVTDSAESREKRASNFVRIGRPSSFVRIGKDVGGDGSFDENDAGQQDYLRDDP
YADEMEKKASSFVRIGRGSNLPLHRFIRIGRMSDYGSYVDDHYNRMYGDEPEAEKRKSHFVRIGKRPSSFVRIGRSGDEVYDDASEDQYDNKRASSFVRI
GKTPVDQLASLKDAIKRASHFVRIGKAPSNFVRIGRNPSSFVRIGRNPSSFVRIGKSELPDDEDFAEKRASSFVRIGKNIGYDLGSSLGYDDLSPADFNS
QDMLKRASSFVRIGKSGDGEDKRASSFVRIGKSGDNSGPHLSVDGTDDGQKRASSFVRIGKSGMDNAADGEGALSSDKRASNFVRIGKALSSPGAPAQPS
SVGSSSSMSSLSSPSSSSSHSDSSSSEESLGDGVLFDDDKPISVASRAHAFVRIGKIPSSAFVRIGKNSDVMVEPALGRRAGYRRGSRGGQSSFVRIGK

*Crassostrea gigas* XP_011448017.3

MLRPYHVIIVGLFYCYTTNAEINENKLLHPIKTEEGANEILGDKADDKRSRGFFRIGKKSAVENEANDKKFDSKTVKEEDNYIPEKIQLIRVVNSESETP
IYVPVEFDPESSDDTVDEDEKRASGFFRIGKSAENVDKRKGFFRIGKSVDQNPMNKKASGFFRIGRTPIDKRGKGFFRIGKSLNEMDEKRASGFFRIGKS
ALNDKRSRGFFRIGRSKGFFRIGKAFPLDGEKRASGFFRIGRNSPEEMRKKASKFFRIGKSVNSKEENDKRASGFFRIGKKCSGDSDKAGDNLTEDKSQS
NPENEDTSESFNRNSDEPVRRASQFFRIGKSSSNKVTKRSSGVNSSPEQNLNLNKRAFFRIGKVPTSAFMRIGRQHLLQSLVSDPLYRNGRIQQSSFIRI
GKRSMSDNHLIDDEQSDSSL

*Octopus bimaculoides* XP_014791297.1

MAGLWRIVLLATITLMCVTSQWVIYVQAEEASSSDELVNSANEADPEMDSEDDLKRANTFLRIGKANGILRLARSPSSFLRIGRRPPLHFVRIGKAPSSM
FLRIGKRVDDSENDDLNEYDADKMSGRDTRASPSSFLRIGKSGNEVEDEIDDTDETVKRVNAFLRIGRQNDPSSFLRIGKSLNNDDLSKDKRTNAFLRIG
KIPASSFIRLGRGPFTEDNGINTRGFRGPTRGFLRIGKRAAIPDGSHADYFSDLNVKSQ

*Lottia gigantea* XP_009051543.1

MECDESSVRRHQPILWSKDSLLSVIDIGQHLHMTTFDEMIQLTRQIMEINLVFSLLLVCSISFVLSHPFTDDSNNDDQDLKSAIQDGDTVPQAVKRPSSF
VRIGRNPSSFVRIGKAFGRFIRIGKNDPNKRLSSFVRIGKSDPNKRISSFVRIGKSQEFNQEPEKRQSSFVRIGKSPELNENEIPNKRYSSFVRIGKSMD
DGSLENPDKRYSSFVRIGKNIENELTNAGLEKRPSSFVRIGKSYFAEPGDMDAEKRLSNFVRIGKSGLEEPEMEQKRAFVRIGKIPSSAFVRIGRMPLYD
AILQKPLGYYNVARRMGKSSFVRIGKRNNEA

*Capitella teleta* 226516

MAQIVHLSLLLVTAASGVLASLTPVCSDLCQKLDVQQLNCVLVCESILVEGPSQVANELDNYDEDDMDKRQKSSFVRIGRNFLPFVRTQRHKSAFVRIGR
STDSLEKRPSSFVRIGRSPYMKRPSSFVRIGKKSDNEEEKRASSFVRIGKAGSKFVRIGKSYDGLDNGEEEQGDRYVRVGEALVNKRPSSFVRIGKKSDE
EEEKRASAFVRIGKALAEKRPSSFVRIGKKSDDEIETMEETPMKRPSSFVRIGKSLAESDGALIPVDEEEKRASSFVRIGKRPSSFVRIGKSLSETSELD
NEKRASSFVRIGKSFPEGQDEELLDQVKRASSFVRIGKRSSDAAE

*Malacoceros fuliginosus* Mfu.C77964.0.T1

MFGCWTLLAGLCVASAVLAMDPELALCEDICSRQGEGEPTECIFLCSQILNGQIDLPITEEPQDEYDEYMPENSLAKRQKSAFVRIGRQKSSFVRIGKKN
AYRDLLKRPSSFVRIGKSYFKRPSSFVRIGRSLAEMEEPAEKRASSFVRIGKSGDASNLYDRDARASSFVRIGKSPSSFVRIGKKASSFVRIGKSSPEEN
EKRASSFVRIGKSSGMDEEKRASSFVRIGKSDPEEKRASSFVRIGKSDESAVPEEKRASSFVRIGKSSGDALEGEDKRASSFVRIGKSLNDDDNQELVNE
KRARSGFVRIGKSYGPSEENEEKRARSGFVRIGKSNSDPIEESAEKRARSGFIRIGKSNSDPIEESAEKRARSGFIRIGKSYNDAIEDNEEKRARSGFVR
IGKSSNDALEDNEEKRARSGFVRIGKSYNDALEDSEEKRARSGFVRIGKSYNDALEDNEEKRARSGFVRIGKSSSDPLDEEKRARSGFVRIGKSYIEPSE
EEKRARSGFVRIGKSYSDGLEENEEKRARSGFVRIGKSSDDEKRARSGFVRIGKSSDGTDEKRARSGFVRIGKSDAQTLDNEKRARSGFVRIGKASSFVR
IGKSLKDEQSKRASSFVRIGKSLQDEDMNSEEKRASSFVRIGKRDAELYQ

*Aplysia californica* NP_001191611.1

MTLHLASPFILLFTIAYSLTSAVQGLEPLPAASLSDSPASGADVPPLPSSAATNAAVDKEWLRQKLEEGQFLPQQDKRWGGINSWMTHRLGGPSERDSSQ
DSLDKQLLVNNVQNYDDSSKRKWSKFSSWGKRDASEETPEGGEGEDGLGAVKKWKNMAVWGKRAEDGLDKRWKQMATWGKREDGDVLGLGTDKRWKQMAS
WGKRLDDSDRDKKWKQMSVWGKREDNGEPLDKKWKEMSVWGKRDTLDDPEKRWKQMAVWGKRQGLDDRNDKRWKQMATWGKRNSSENYDKRWKQMSVWGK
RDGDGDLDKRWKQMSVWGKRDGDGDLDKRWKQMSVWGKRNGDGDLDKRWKQMSVWGKRDGDEDVEKRWKQMSVWGKRDGDADLDKRWKQMSVWGKRDEDG
NLDKRWKQMSVWGKRDEDGNLDKRWKQMSVWGKRDGDDNLDKRWKQMSVWEREREREWV

*Capitella teleta* 197880

MASCRLLLTVITLVICSLVVLADDPETEEQAQDLVPHDMDQDLMDKRKWGSNSMRVWGKRDGDDEMEMDGGAEKRKWGGNNNMRVWGKRKWGANSMRVWG
KRSELPEEEKRKWGGSNTMRTWGKRADDNEEDELAKRKWGSNSMRVWGKRADDNKRKWGSNSMRVWGKRADDDMDESKRGWKNNNMRVWGKRADDEIDED
KRKWGSNSMRVWGKRSADDDAELAAAVPHAIVKRSLDSEEFTDDMEKRRWGGNDMRVWGKRRSRADGPKRSWKTNVMRVWGKRGWADNNMRVWGKRADEG
AEKRAWVGDKSLSWGKRSDNEVIRNLLAEQVMMISIISPTKYLRDVAICVGGVLGGFSRDPLSE

*Malacoceros fuliginosus* Mfu.C51764.1.T1

MKACPLLLALAACVLVVVSAKEDPKPQPDAATPQLKTTFPSSISHNSNDVIAKPNQNIDETADGDNEGNELEKRQWNNLRGSMTWGKRGWNDNSRLWGKR
ADENEVESIKDEDQKRKWGEGSRLWGKRSDDMDKRKWGSNRLWGKREEGHKKWSSNMNMWGKRSDSEVTDHEEEKRKWGDSSRLWGKRSEDDSEMDEEKR
KWGGNSRMWGKRSVSDEDVKRKWGGNSRMWGKRDDELTEEDKRKWGGNSRMWGKRSDVEEPEEKRKWGGNSRMWGKRSDELTEEDKRKWGGNSRMWGKRA
DDQDMEEEKRKWGGNSRMWGKRMSELPEEEKRKWGGNSRMWGKRSDDEEMDEEKRKWGGNSRLWGKRSDEMIEEAKRKWGGNSRMWGKRDGDNDLLLDEE
EKRKWGSARTWGKRSEADDNDDKRAWKSQGSRLWGKRSKRSIDDFLEPAEKRAWYSKSNRLWGKRTNPYSTSYGGPKRGWKANSMRVWGKRSSWKNMNMW
GKRDALADDEADDQESYAELMNSLNDYIDNVKRAWHSGSSRVWGKRSIDDTMDSDSAL

**Figure S2. Selected pro-peptides.** Putative neuropeptides, based on double basic residue (e.g. KR) and GR cleavage and amidation sites, are highlighted in color within the pro-peptide: blue, FMRFa; orange, LFRYa; green, FVRIa; magenta, Wamide (or "myoinhibitory peptide"). All sequences/accession codes from NCBI, except for *Capitella teleta*, from Capcal1 genome, Joint Genome Institute, and *Malacoceros fuliginosus*, from our transcriptome resource (38).

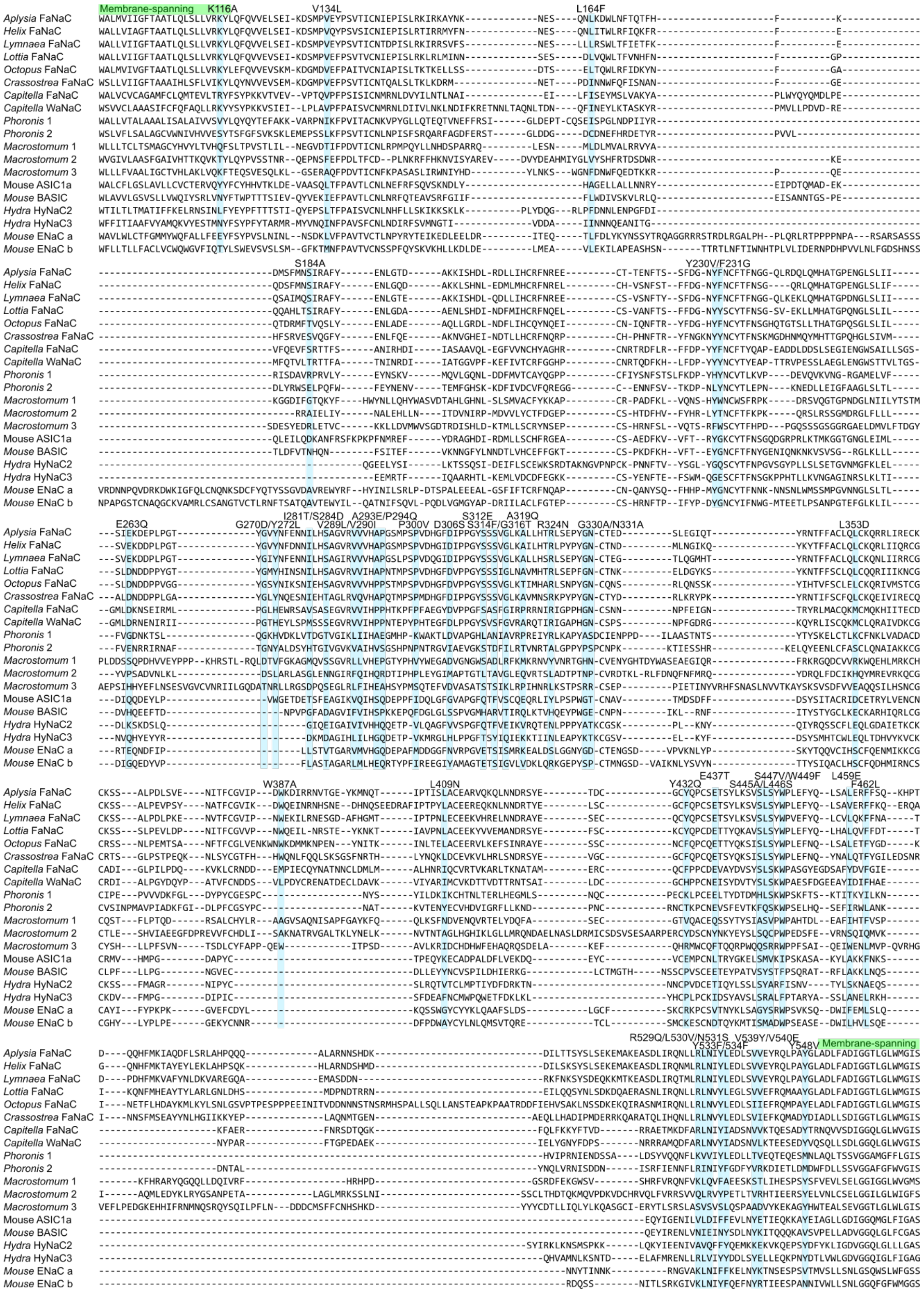


**Figure S3. FaNaC-specific amino acid residues.** Alignment highlighting 43 positions where channels gated by FMRFa possess physico-chemically similar residues and non-FMRFa-gated channels possess different residues. 30 *Aplysia* FaNaC mutants based on these differences are indicated above the alignment. Putative intracellular domains (before and after green membrane-spanning domains) have been excluded.

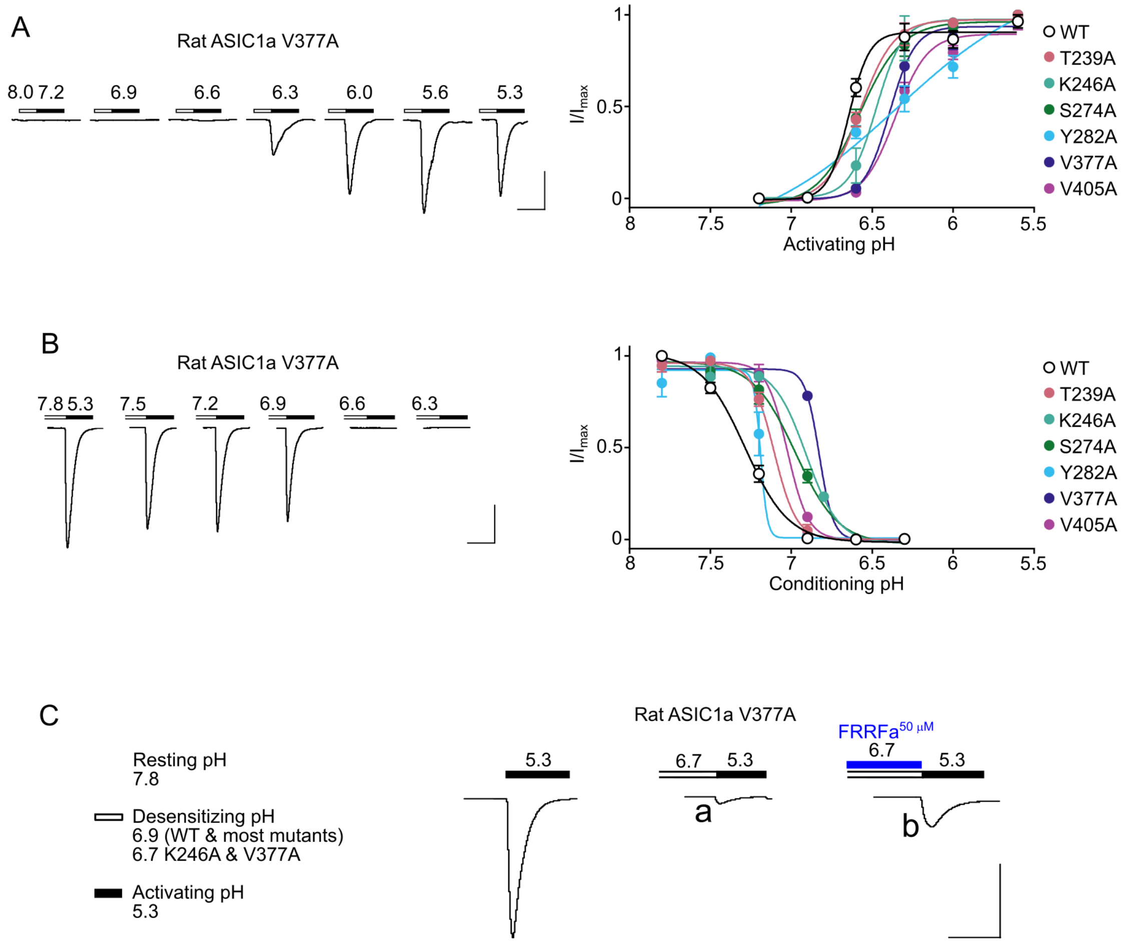


**Figure S4. Measurement of steady state desensitization and peptide modulation of rat ASIC1a mutants.** (**A**) *Left*, example currents at an oocyte expressing a rat ASIC1a mutant in response to decreasing pH. *Right*, mean (± SEM, n = 3-4) normalized currents in response to decreasing pH. (**B**) *Left*, example currents at an oocyte expressing a rat ASIC1a mutant in response to pH 5.3 after pre-incubation in decreasing pH. *Right*, mean (± SEM, n = 3-4) normalized currents in response to decreasing pre-incubation pH. This established that pH 6.9 was strongly desensitizing for WT and most mutants, and pH 6.7 was strongly desensitizing for mutants K246A and V377A. (**C**) Example experiment showing: current response to activating pH (5.3) after pre-incubation in resting pH 7.8; current response ("a") to activating pH after 20 s pre-incubation in desensitizing pH (6.7 for this mutant); and current response ("b") to activating pH after 20 s pre-incubation in desensitizing pH plus 50 µM FRRFa. Dividing current amplitude "b" by that of "a" gives fold-enhancement by FRRFa in Fig. 6B. All scale bars: x, 10 s; y, 10 µA.

### Supporting text

**Oocyte expression vector and novel cDNA inserts.** For oocyte expression of previously uncharacterized genes, we modified the pSP64poly(A) vector (ProMega) to include 5' and 3' untranslated regions (UTRs) from the *Xenopus laevis* β-globin gene, as described elsewhere (41), and commercially synthesized FaNaC inserts (Genscript) were sub-cloned between SalI and BamHI sites, except for *Lottia* FaNaC_XM_009055314.1, which was subcloned between these sites in the unmodified pSP64poly(A) vector. Sources/databases for these accession codes are described in *Experimental procedures*.

```
Sp6, HindIII, 5'UTR, SalI, Kozak consensus sequence, XbaI, BamHI, Myc + TAA stop codon, 3'UTR, poly(A), EcoRI

>Modified_pSP64_vector
ATTTAGGTGACACTATAGAATACAAGCTTGCTTGTTCTTTTTGCAGAAGCTCAGAATAAACGCTCAACTTTGGCGTCGACTCTAGAGGATCCGAGCAGAAGCTCATCAGTGAGGAAGATCTCTAAGGT
TACCACTAAACCAGCCTCAAGAACACCCGAATGGAGTCTCTAAGCTACATAATACCAACTTACACTTTACAAAATGTTGTCCCCCAAAATGTAGCCATTCGTATCTGCTCCTAATAAAAAGAAAGTTT
CTTCACATTCTAAAAAAAAAAAAAAAAAAAAAAAAAAAAAAACGAATTCGTAATCATGTCATAGCTGTTTCCTGTGTGAAATTGTTATCCGCTCACAATTCCACACAACATACGAGCCGGAAGCATAAA
GTGTAAAGCCTGGGGTGCCTAATGAGTGAGCTAACTCACATTAATTGCGTTGCGCTCACTGCCCGCTTTCCAGTCGGGAAACCTGTCGTGCCAGCTGCATTAATGAATCGGCCAACGCGCGGGGAGAG
GCGGTTTGCGTATTGGGCGCTCTTCCGCTTCCTCGCTCACTGACTCGCTGCGCTCGGTCGTTCGGCTGCGGCGAGCGGTATCAGCTCACTCAAAGGCGGTAATACGGTTATCCACAGAATCAGGGGAT
AACGCAGGAAAGAACATGTGAGCAAAAGGCCAGCAAAAGGCCAGGAACCGTAAAAAGGCCGCGTTGCTGGCGTTTTTCCATAGGCTCCGCCCCCCTGACGAGCATCACAAAAATCGACGCTCAAGTCA
GAGGTGGCGAAACCCGACAGGACTATAAAGATACCAGGCGTTTCCCCCTGGAAGCTCCCTCGTGCGCTCTCCTGTTCCGACCCTGCCGCTTACCGGATACCTGTCCGCCTTTCTCCCTTCGGGAAGCG
TGGCGCTTTCTCATAGCTCACGCTGTAGGTATCTCAGTTCGGTGTAGGTCGTTCGCTCCAAGCTGGGCTGTGTGCACGAACCCCCCGTTCAGCCCGACCGCTGCGCCTTATCCGGTAACTATCGTCTT
GAGTCCAACCCGGTAAGACACGACTTATCGCCACTGGCAGCAGCCACTGGTAACAGGATTAGCAGAGCGAGGTATGTAGGCGGTGCTACAGAGTTCTTGAAGTGGTGGCCTAACTACGGCTACACTAG
AAGAACAGTATTTGGTATCTGCGCTCTGCTGAAGCCAGTTACCTTCGGAAAAAGAGTTGGTAGCTCTTGATCCGGCAAACAAACCACCGCTGGTAGCGGTGGTTTTTTTGTTTGCAAGCAGCAGATTA
CGCGCAGAAAAAAAGGATCTCAAGAAGATCCTTTGATCTTTTCTACGGGGTCTGACGCTCAGTGGAACGAAAACTCACGTTAAGGGATTTTGGTCATGAGATTATCAAAAAGGATCTTCACCTAGATC
CTTTTAAATTAAAAATGAAGTTTTAAATCAATCTAAAGTATATATGAGTAAACTTGGTCTGACAGTTACCAATGCTTAATCAGTGAGGCACCTATCTCAGCGATCTGTCTATTTCGTTCATCCATAGT
TGCCTGACTCCCCGTCGTGTAGATAACTACGATACGGGAGGGCTTACCATCTGGCCCCAGTGCTGCAATGATACCGCGAGACCCACGCTCACCGGCTCCAGATTTATCAGCAATAAACCAGCCAGCCG
GAAGGGCCGAGCGCAGAAGTGGTCCTGCAACTTTATCCGCCTCCATCCAGTCTATTAATTGTTGCCGGGAAGCTAGAGTAAGTAGTTCGCCAGTTAATAGTTTGCGCAACGTTGTTGCCATTGCTACA
GGCATCGTGGTGTCACGCTCGTCGTTTGGTATGGCTTCATTCAGCTCCGGTTCCCAACGATCAAGGCGAGTTACATGATCCCCCATGTTGTGCAAAAAAGCGGTTAGCTCCTTCGGTCCTCCGATCGT
TGTCAGAAGTAAGTTGGCCGCAGTGTTATCACTCATGGTTATGGCAGCACTGCATAATTCTCTTACTGTCATGCCATCCGTAAGATGCTTTTCTGTGACTGGTGAGTACTCAACCAAGTCATTCTGAG
AATAGTGTATGCGGCGACCGAGTTGCTCTTGCCCGGCGTCAATACGGGATAATACCGCGCCACATAGCAGAACTTTAAAAGTGCTCATCATTGGAAAACGTTCTTCGGGGCGAAAACTCTCAAGGATC
TTACCGCTGTTGAGATCCAGTTCGATGTAACCCACTCGTGCACCCAACTGATCTTCAGCATCTTTTACTTTCACCAGCGTTTCTGGGTGAGCAAAAACAGGAAGGCAAAATGCCGCAAAAAAGGGAAT
AAGGGCGACACGGAAATGTTGAATACTCATACTCTTCCTTTTTCAATATTATTGAAGCATTTATCAGGGTTATTGTCTCATGAGCGGATACATATTTGAATGTATTTAGAAAAATAAACAAATAGGGG
TTCCGCGCACATTTCCCCGAAAAGTGCCACCTGACGTCTAAGAAACCATTATTATCATGACATTAACCTATAAAAATAGGCGTATCACGAGGCCCTTTCGTCTCGCGCGTTTCGGTGATGACGGTGAA
AACCTCTGACACATGCAGCTCCCGGAGACGGTCACAGCTTGTCTGTAAGCGGATGCCGGGAGCAGACAAGCCCGTCAGGGCGCGTCAGCGGGTGTTGGCGGGTGTCGGGGCTGGCTTAACTATGCGGC
ATCAGAGCAGATTGTACTGAGAGTGCACCATTCGACGCTCTCCCTTATGCGACTCCTGCATTAGGAAGCAGCCCAGTAGTAGGTTGAGGCCGTTGAGCACCGCCGCCGCAAGGAATGGTGCATGCAAG
GAGATGGCGCCCAACAGTCCCCCGGCCACGGGGCCTGCCACCATACCCACGCCGAAACAAGCGCTCATGAGCCCGAAGTGGCGAGCCCGATCTTCCCCATCGGTGATGTCGGCGATATAGGCGCCAGC
AACCGCACCTGTGGCGCCGGTGATGCCGGCCACGATGCGTCCGGCGTAGAGGATCTGGCTAGCGATGACCCTGCTGATTGGTTCGCTGACCATTTCCGGGTGCGGGACGGCGTTACCAGAAACTCAGA
AGGTTCGTCCAACCAAACCGACTCTGACGGCAGTTTACGAGAGAGATGATAGGGTCTGCTTCAGTAAGCCAGATGCTACACAATTAGGCTTGTACATATTGTCGTTAGAACGCGGCTACAATTAATAC
ATAACCTTATGTATCATACACATACG
>Crassostrea_FaNaC_XM_011442205.2
GTCGACACCATGAATGGATATATAGCAAATGGGGGTCAGAATAAAGAACACATGAAGTTTTCACCTGTAGGAAAGTTTAACTATTTTCACGACCATTTGAAGATGCCGCCAAATGACACCCATGCTCG
AAAGAAAAATCGAAAATCGGCCATGGCCATAGTGCAGGAACTGGGGTCCGAAAGTAATGCACATGGTCTAGCGAAAATCGCCATGTCACGAAAAACTAAACGAAAAGTGATGTGGTCTCTGCTGGTGA
TAATTGGGTTTACGGCAGCCGCCATACATTTGTCATTCCTTGTGATAAAGTACCTACAGTACAATGTGGTGGAAGTCTCCGAAATGAAGGACGGAATGCCGGTGGAATTTCCATCGGTAACAATCTGC
AACACGCAGGCTTTATCGCTTACCAAACTGAAGGACAGAATGAATGAAACTCCCGATATCAACAACTGGTTTCAATTTATTAGCAATGCAAACTTTGGTGAACACTTCTCGCGAGTGGAGTCTGTTCA
GGGATTTTACGAGAATTTATTCCAGGAAGCCAAAAACGTGGGCCACGAAATAAACGATACGTTACTTCACTGCCGATTCAATCAAAGGCCGTGTCACCCTCACAACTTCACTAGATATTTCAACGGCA
AGAACTACTATAACTGCTACACGTTTAACAGTAAGATGGGGGACCATAACATGCAGTATATGCATACCACCGGTCCCCAGCATGGCCTCTCAATCGTCATGGCACTGGACAATGACGACCCTCCGCTA
GGGGCATACGGACTCTACAACCAAGAGAGTAACATTGAGCACACCGCTGGACTGAGGGTCCAGGTTCATGCCCCCCAGACGATGCCCAGCCCCATGGATCACGGATTTGACATCCCTCCTGGGTATTC
GTCCTCCGTGGGTCTAAAAGCCGTGATGAACTCAAGGAAACCTTATCCTTATGGAAACTGTACATATGACAGACTTAAGAGATACCCAAAATACAGGAATACGATATTCTCCTGTTTCCAGTTGTGTA
AGCAAGAGATAGTGATTCGCGAGTGTCAGTGTCGGACCTCCGGACTCCCAAGCACTCCGGAACAGAAAAATCTGTCTTACTGCGGGACATTCCATCACTGGCAGAACTTATTCCAACAGCTAAGTAAG
TCTGGCTCATTCAATAGAACACACCTGTACAACCAGAAACTGGACTGTGAGGTCAAGGTGCTCCACCGTCTCTCTAACGACCGCAGTTACGAGGTCGGGTGTGGGTGTTTCCAGCCATGTCAAGAAAC
CTCCTATCAAAAATCCATCTCCCTCTCGTACTGGCCCCTCGAGTTTAACCAGTACAACGCCTTACAGACATTTTATGGAATTTTAGAAGATTCAAATCGGATAAATAATTCCTTCATGTCAGAGGCTT
ATTACAATCTTCATGGAATTATTAAGAAATATGAGCCACTGGCTCAAAACATGACTGGAGAAAATGCGGAACAATTATTACATGCCGATATTCCGATGGATGAGAGACGAAAGCAGGCCCGTGCCACT
CAACTCATTCATCAAAATCTCCTTCGTTTGAATGTTTATTTAGAAGACCTTAGTGTCATAGAATTTAAGCAGATGGCTGACTATGATATAGCAGATTTACTGTCAGATATCGGAGGAACTTTAGGACT
GTGGATGGGGATTTCGATTCTGACAATTATGGAGTTAGTGGAACTCGCGATTCGACTGGTTGCCATTTTATTCAAATCGGAAAATAGATATCCTGATCACATGGATGACCACAGTTCAGCCAATGGAA
TTTTAGAACACTCGACAGAGAGGGCGATATACCCCCAAGATACATATGATCCATACACACAGCCTGAGTTTCAGAGTTTTGAAAAAAATGATTATGGTCGGACTACAAACGATTTACCTCCCGACTCG
CCTATCGGATCC
>Octopus_FaNaC_XM_014930938.1
GTCGACACCATGAAGGTAACAGGCTTTGACAAACTGGGTTTCAACAGCAACCAGTTACGCACAATGAGGCAACAGCTACGCAGGCGGAACTATAATGCGCTCAGCATCATCACAGAATTGGCAGCCGA
GAGCAATGCCCACGGACTGGCAAAAATTGCAACATCAACACAGACCCCAAGAAAAGTTCTGTGGGCATTAATGGTAATTGTAGGCTTCACTGCCGCCACACTGCAACTCTCTCTTCTCGTCCGGAAAT
ACCTGGAGTTCCAGGTGGTAGAAGTTTCTAAAATGAAAGATGGTATGGATGTGGAATTTCCTGCCATCACTGTTTGCAATATAGCACCAATATCATTGACGAAAACCAAAGAGCTGCTTAGTTCAGAC
ACGTCAGAACTGACACAATGGCTGCGATTTATAGACAAATATAACTTTGGTGCACAGACTGACCGGATGTTCACGGTGCAGAGTTTGTACGAGAACCTCGCAGACGAGGCCCAGCTTTTGGGTAGGGA
CCTCAATGATTTCCTCATACACTGTCAGTATAATCAGGAAATCTGTAACATCCAAAACTTTACTCGATATTTCGACGGACATTATTTTAACTGTTACACATTCAACAGCGGCCATCAGACCGGCACCA
GCCTGTTGACACATGCCACTGGGCCTCAGAGTGGTCTTTCTTTAATTCTTTCGTTAGACAATGATGATCCACCTGTTGGTGGCTATGGCTCATACAATATAAAATCAAACATTGAACACAGTGCTGGA
GTAAGAGTGGTTGTTCACCCACCAAGCACGATGCCGAGCCCAGTAGACCATGGCTTTGACGTGCCACCAGGGTACTCTTCATCAGTAGGTTTGAAAACTATTATGCACGCACGTCTTTCAAATCCCTA
TGGCAACTGCCAAAATAGCCGGCTTCAAAATTCCTCAAAATATATCCACACCGTTTTCTCATGTCTTGAACTGTGTAAACAGCGAATTGTGATGAGCACATGCGGCTGTCGATCTTCAAATTTACCAG
AAATGACATCGGCAAACTTCACTTTCTGTGGTTTAGTGGAGAACAAATGGAATTGGAAAGATATGATGAAGAATCCAGAAAACTACAACATCACAAAAATCAACTTGACAGAACTTGCCTGTGAAGAA
AGAGTTCTCAAAGAATTTTCCATAAACCGAGCTTACGAAAGCTCATGTAATTGTTTTCAACCATGTCAAGAGACCACATACCAAAAATCAATATCTCTTTCCTATTGGCCTTTAGAGTTCAACCAGTT
AAGTGCCCTTGAAACATTTTATGGCGATAAAATCAATGAGACATTTTTACATGATGCTTACAAGATGCTTAAATACTTGTCAAATCTTGGTAGTGTCCCCACACCTGAGTCCCCACCTCCGGAAGAGA
TCAATATCACTGTGGACGATAACAACAACAGCACCAACTCTAGGATGCATTCTCCGGCTTTACTGTCGCAGTTGCTGGCAAACAGCACTGAGGCACCAAAGCCTGCAGCAACGAGAGACGACTTCATT
GAACACGTGAGTGCAAAGCTCAACTCAAGCGACAAAGAGAAACAAATCCGAGCGTCAAACATGATCCGACAGAACCTTCTGCGACTGAACGTTTACCTTGAAGACCTCAGTATCATTGAGTTCCGACA
GATGCCTGCCTACGAGCTGGCTGATCTGTTTGCCGACATTGGCGGCACACTGGGTCTGTGGATGGGTATATCGGTGCTGACTATTATGGAGCTTATAGAACTCTTCACCAGACTTCTAATGCTAATTT
TCAGCTCGGAAAAGAAAATTCCCAATGCCGACCCAGTCACAAATGGAATGTTGGACCATGACTGTGACTGCCAAAAAAGTAGTATGGAATCACCATTTGGATCC
>Lottia_FaNaC_XM_009055314.1
AAGCTTACCATGGTGCATCGAAACACGGGAAAAAATCAAACAGTACTCTCCCTTGTGGCGGAGCTGGGATCGGAAAGTAACGCTCATGGATTAGCGAGAGTAGTGACATCACGAGAGACCAAACGGAA
AGTGATCTGGGCCTTATTAGTGATTATCGGATTTACTGCTGCTACGTTACAATTATCATTACTGGTGAGAAAATATCTTCAGTTTCAAGTCGTAGAGGTGTCAGAGATTAAAGACAGCATGCCTGTCG
AGTTTCCCTCGGTGACCGTGTGCAATATTGAACCCATTTCTTTACGGAAATTACGTCTAATGATAAACAATTCTGAAAGTGATTTAGTGCAGTGGTTAACGTTCGTCAACCATTTTAATTTCGGTCCG
CAGCAAGCCCACTTAACATCGATCCGCGCTTTTTATGAAAATCTGGGCGATGCAGCAGAAAATTTAAGCCACGATATCAACGATTTTATGATTCATTGTCGGTTTAACCAAGAATTGTGTTCCGTGGC
CAATTTCACCAGTTTTTTTGACGGAAATTATTATAGTTGTTACACATTTAATAGTGGCTCTGTGGAAAAAATTACTCATGCACGCCACTGGTCCTCAAAATGGCCTCTCGTTAATTCTTTCTTTGGATA
ACGACGACCCTCCATATGGAACATACGGAATGTATCATATTAATTCTAATATTCTACACAGTGCTGGAGTCCGCGTAGTGATACATGCACCTAACACCATGCCTAGTCCCGTGGATCACGGTTTCGAT
GTGCCGCCAGGATATTCATCTTCGATTGGATTAAACGCTGTGATGCATACGCGTCTTTCAGAACCGTTTGGAAATTGTACAAATAAGGAGTTAGATGGATACAAGAGTTACAAGAACACATTTTTTTC
TTGTCTTCAATTGTGTCAACAACGGATTATTATCAAAAATTGTGGATGTAAATCGTCCAGTTTACCAGAAGTGTTGGACCCAAACATCACATTTTGTGGAGTCGTTCCCAACTGGCAAGAAATTCTAA
ATCGTTCGACTGAATACAAAAACAAAACTATTGCTGTTCCAAATTTGGCATGCGAAGAAAAATATGTGGTCGAAATGGCCAACGATCGAAGTTACGAATTTTCGTGTAAATGTTACCAGCCATGTGAC
GAAACGACATATCAAAAAGCCGTTTCTCTCTCATACTGGCCGTTGGAGTTCTACCAACTTCACGCATTACAAGTCTTCTTCGACACAACCATCAAACAAAACTTTATGCACGAGGCATACACTTACTT
AGCGCGTCTGGGCAACTTGGACCATTCAATGGACCCCAACGATACCAGACGCAACGAAATTTTACAACAAAGTTACAATCTAAGTGACAAAGACCAAGCAGAGCGAGCGTCAAATTTAATCAGACAAA
ACCTGCTGAGACTCAATGTATATTTGGAGGGGTTAAGTGTGGTGGAGTTCCGTCAAATGGCGGCATACGGATTAGCTGATTTATTCGCAGACATTGGTGGTACTTTGGGACTGTGGATGGGTATCTCA
GTTTTGACAATCATGGAACTGATAGAACTAATAGTCAGACTTTTTTTGCTTCTGTTTAACTCCGAGAAGAAGATCCCAGAACAAGATCCTGTATCCAATGGCATGCTAGAAACTGAACGAGATTACGA
GTATGAACGAACTGGTGTGGAATCGCCGGTCGGATCC
>Pinctada_construct_pfu_aug2.0_4392.1_09318.t1
GTCGACACCATGGATTCCGACGTTGTAAATGAGGGACCAACGCCAAAGAAACGGCAACTAGTACGGATGCGAACAAGGGATGTAATTCACAAACTTGGTTCAGAGAGCTATATGCATGGGATTTCGAA
GATAATGGCTTCCAATTCATTTAACAGGAGATTGTTTTGGACTATTCTTTTACTATGTGGAGTTATAGCTGCAGTAACACAGCTATCCATCTTGTCTTTAAAATATTTGAAATTCGACACATTTTTTA
```

TTACAACAGAAATACAGAACTCGGAGGTAGAATTTCCATCTATCACTGTATGCCCGAATTATGTTCCCTTCAAATATCCATCCGAGGTACTTAAGGACACAGAAATTCAACTTTACGAGGACTGGTTA
AGTTTTCTTCTTAAGTTAGAATATCCAAGTACAGAGTACGACCATTTTAGTTCATGGACCGGCTACTATGAAAATCTCCCCGATATTCTACCCCTTTTCGCCACCGAGTGTAGAGATATGATTTTGCA
ATGTACTTATCGAGGGAAAAGTTGTTCTTGTGAACACTTTGATGAGTATATAAATGCTGAGTATGTTCTTAAATGTTATACTTTCAACGCATCTAAATTTATGACGATACCAAGCATGAAATATGCAA
CATCTGGACCAGATCTAGGATTGTCGTTGATCCTAGGAGTGGATGATACAAAGTTTGAGGACAAATTTAACATTGTAGGGGATACTTCAAGACCTTATGATCTCGATTACTTGTACGGCTATAGTTCC
GGGTTCACAGTTCAGATACACCCTCCCGATACAATCCCAACACCAATGACACATGGTTTCGACATATCAACAAGTCAATCAACAACAGTGGCACTAAAAGAATACATAGAAGGTTATCTACCACATCC
ATACACTAATTGTTCATTTGATGCATTGCGAAGTGATAAAATATACCGTCGAACATTGTTCACATGTGACCTAATATGTAAGCAAAAGAACGTAATCAAGAACTGTGGATGTACCTCCTCGATTCTGC
CTAATATTGGTAAATACCAGATCTGTGGACAATTGAAAAATTGGAATAGCGATCCTAACATAACAGATATCATACAAAATTTGTTGTGCGAGAAAAAGAACATTCTAAATAATACGCCATGTCATTGT
TCTCCATGTAAAGAAAATACGTATGATATTACATTATCGGCTTCCGATTGGCCAAAAGACGGTCATGTATTCAAACTACTTGAAGGCCTTCAAGAATATTTAAACGAAAACTTGCGTATGGGATATGA
TAAATTACGCGATGAAATGTCAGACACTCAAAATTTTGAAAAAATAAATTCTTACATACGGAAAGCAAAGTTAGATTTGGCAAGACTAAATGTTTACTTTAAAGAACTTAATGTTGTGTCAACAGTGC
AGATTCCAAGTTATCAATTCGCCAATTTAATGGCCGATATAGGAGGATCACTTGGTCTTTGGATTGGAATTTCTGCTTTGTCCATGATGGAGCTATTAGAACTATTAACATTAATTATGTGTGCATTT
AGAAAAAAGAAGAAAAACTACAATGCTTCAAAACAAGACGACGGATCC
>Capitella_Capca1_207658
GTCGACACCATGAAGAAACAGGACAGCAGCCAAGAACTGAGCACGGCTCAGAGGATGAAAGAATCTCTACTGACGTTCTGCCGTCACACTACGGCGCATGGCTTGTGTACTGTGCCGTATTCGTCTAA
TAAATTCAGTCGCGCTATATGGCTGACACTGTTTTTTGCAGCTTTGGCCGGAATAGCCTATCAGACAACAGGAATAATGACAGCATATTATTCATACCCAGTACAAGAAGTCAACCATATGGACCAGG
GTCCAGTTCCCTTTCCAGACGTCACAGTGTGCAACATCGAACCTATAGTGATGAGCAAAGCTCTTGAATTAGCCAATGATCCATCAACAGCCACGCATGAATATAACGAGAAGAAGGTTCCTCTCATA
CGGGACCTCTTGTTGAGAAATCCGACGTCGATTGCGGGCATGGCCAGCAGAGTTGGTACCTCCAAAGGCTACTTTGAAAACCAACCCCAGAACGAGAGCCGCTACGCCAGTCACTCTCTGAAGGACCT
CATCATAGGCTGCTCGTACATACACAAGAACTGTGTGGGAAATCTGGAGGAGAGTTTCACCTACAAAGCGGATGGCACCTATTTCAACTGCTACACCTTCTCTCCAGCCAATAACCCCAACCTATCCG
ATAACACCGTGGGCCTTGGACCTGAATCTGGCCTCTCTCTTATCCTCTTCATGGATACTGACACAGCAGACCAAACCTTCCCGGGGATAATAGATGTCCAGTCGAATGTGAAGCAAAGCGCGGGTGCC
CGGGTGACCGTCCACAATAGAGGAACACTTCCTAACCCCCTCATAGAAGGTTTTGACATAATGCCTGGTCATTCGACGTCAGTCGGAGTGACTGCTACTCGGATACGGCGCCTCTCGGAGCCGTACAC
TAACTGTACGGACGGAGCTTGGCTGAAATTGAAACCGCACTTTGAGTACACTCCGACTTCTTGCTTGCTGCTCAGCGCCCAAGAATTTATATTCAAGAATTGTGGATGCCTGTCCGCTGAACTTCCTG
TACCCGACAATCTGTTTGGTGAGAAATACTGCGGATTTTACGATTATGACAACAACACCTACCTAGAACAAATCGAATGTGAACCCAAGGCTTATGTTAATTTCTACACTGCTACTGATGCCGTCAAG
CCAAACACACACTGCCGACCGCTCTGTGATACGTTCCAGTATGATTTCCTTCTGTCGGAAGCATACTGGCCAAAGGAGCGATACGAAGCCAGCTTCTTTGAGGAGTACGTTGCCAGTCGCCCCGATCG
AGACTCTCTGCAGGCTTACACATCCCTTGTGAACTACTCAACGACTGAAGACTCAAAGATCAGAAGAAACTTTGCTCGAGTCAACATTTATATGAAGTCCTTAGAAACATTGACTAGATCGCAGATGC
CGTCCTATCAAAGCTCCAACTTGCTGTCGGACATCGGTGGAACATTGGGTCTTTGGGCTGGCATCTCTATCATCACCGTTTGTGAGATCATTTCTTTTGCCGTGCGCCTTGCAGTCACTCTGCCGCGA
AAGGAAAATATCGGATCC
>Capitella_FaNaC_Capca1_52833_alt_spl_var_Capca1_191096
GTCGACACCATGGACATCAGTATAGAAAAGCAATCGTACACTGCACAAAACCTTTTTACAACAACTGGGCAGCACTGCAAAGTACCTACTGTGCTTGAGGAATTUGCCACTCGGACCACAATGCATGG
TGTTCCAAAAGTGATCAAAGCCAAGTCTTCCTTAGCTCGCATCTTCTGGGCCCTGGTCTGTGTATGCGCCGGGGCAATGTTCTGTCTGCAGATGACAGAAGTACTCACCAGGTACTTCTCCTATCCCA
AAAAGGTGACAGTGGAAGTGGTCCCCACCCAAGTTCCCTTCCCCTCAATTTCGATCTGCAACATGAGGAATCTGGACGTTTACATCTTAAACACTTTAAATGCGATGTTTATACAGAACGACAACCCA
ATCAACAACGTCAACAAAAGTGAGATCCTTTTCATCTCGGAGTACATGTCGCTCGTAGCCAAGTACGCTCCGTTGTGGTACCAATACCAGATGGACCTACCGGAGGTATTTCAAGAAGTATTCTCACG
AACTACCTTCTCCGCCAACATACGACATGACATCATAGCCTCGGCAGCCGTCCAACTGGAGGGATTTGTAGTGAACTGCCATTACGCGGGACACCGTTGCAACCGGACGCGGGATTTCCTCCGCTTCT
TCGATCCGTATTATTTCAACTGCTTCACGTACCAAGCACCAGAAGCAGATGACCTGGATGATTCCCTGAGTGAGGGTATTGAGAACGGTTGGTCCGCGATTCTGCTATCCGGATCAGGAATGCTGGAT
AAGAACAGTGAGATCCGGATGTTGCCTGGACTTCATGAGTGGCGATCCGCAGTCTCTGCCAGTGAGGGTGTCCGCGTGGTCATTCATCCGCCGCACACGAAACCTTTCCCCTTTGCGGAAGGATATGA
TGTTCCGCCTGGATTCTCCGCTTCGTTCGGGATTCGCCCGCGACGGAATATACGAATCGGACCACCACATGGGAATTGCTCGAATAACAACCCCTTTGAAATTGGGAATACGCGTTATCGCCTAATGG
CATGCCAAAAAATGTGCATGCAGAAGCACATCATCACAGAGTGCGACTGTGCTGATATTGGTTTGCCAATTCTGCCAGATCAGAAGGTTAAACTTTGTCGTAATGATGATGAAATGCCCATTGAGTGT
CAGTACAATGCCACCAATAACTGCTTAGACATGTTGATGGCTCTGCATAACAGGATACAATGTGTGAGAACTGTGAAAGCTAGACTCACCAAGAACGCTACGGCGATGGAGCGATGCCAGTGCTTTCC
ACCTTGTGATGAGGTAGCTTACGACGTCTCCTACAGCCTTTCCAAGTGGCCCGCGTCTGGCTACGAAGGTGATTCCGCCTTCTACGATGTGTTCGGCATTGAGAAATTTGCGGAGAGATTCAACCGGA
GTGATACGCAAGGAAAGTTTCAGCTATTCAAGAAGTATTTCACGGTTGACCGTCGTGCGGAAACCATGAAAGACTTTGCCCGACTCAATATTTATATCGCGGACTCCAACGTGGTGAAAACGCAAGAG
TCCGCAGACTACACGAGGAACCAGGTCGTTTCAGACATTGGTGGCCAGCTTGGTTTGTGGGTGGGAATATCCATTATAACCCTTGCTGAAGTTCTTGAACTTATGCTGGATATTCTCCGTTATTTTAC
GTCATCGTCATACATCACGGTATCGACCAGTCCGATGGTCAACAATAACCAAAGCAGAGATATTTCCAATCGTTACAGCATGCCAGAGACTAAATTGGGATCC
>Capitella_Capca1_185559
GTCGACACCATGCCTGGGTCCTCTCGGGGACGCCGCTCCCTCTCCCTCGGGGACGTACGTCACCATGACGACCGAGTCAGCGACGTGCACCGAGTTTTCCAGGAGTTCGCGGACGGCACGTCAATGCA
CGGCGTGCCCCGCATCATCAACGCCCGTTCGATCCCCGGCCGCTTCTGCTGGTCTGTGATCTGCATGGGCGCCTTTGGGATGTTCCTCTGGCAGTGTGGCATTCTCCTGGAACGCTATTACTCATACC
CGAAGAAGGTGATGGTGGAGATTGTCCAGCGACCCGTGCCATTCCCAGCAGTTTCGTTATGTAATACAGACTTCCTGGACCTGGAAGTCGCTCACCATTTGAAGGAAACTCTCATACAAAGCAACGGG
ACATTTATGCCAACGGAAAAACAAGCCAAGTTCATCGAGAGATACTTCGATTTCTTTCTGTACTCGTTCCATTTACTGTCAGTTTATCAAGAGATAACTCAAGATAAGAATAAGTTACAACAGGACAT
GGCTGAGGTATATTCCAGGTTGGGATTGGTCGCGAACCTCGGCCACCAGTTGTCCTCAACGGGCAGCATCCAACTGGACAACTTCATCGTCAGCTGCCGTTTCATGGATGGATTATGCAATGTGACGA
CTGCGTTTGATAAGATATTTGACCCATATTTCTTCAATTGCTTCACTTTCCAACCAAACACAATTCTAAAGTCGAGAGCTTCCAGGCTTCAAGGCGTGGAGTACGGACTGACGGTGCTCCTCTTCACC
AAGAGCGCTGGTCAGGTCGGTATGGATAGGGAGGACGCAGAGGAGTTGACTTTTATTCCTGGGCTCCAAGAGTCTGATTCAGCTCTGGCCAGTGGTCAAGGGGTCCGACTGGTCATCCATCCACCTAA
CACCAGGCCCCACCTAACCGCTGACGGCTACGATATTCCACCAGGATTCTCCGTGACGATTGGCGTGAAAGCGCGCGAAAATGTGCGCATTCGGCATCCTCATGGCAACTGCTCATCCATGAAAAATG
TACCACACGTAGAGGAAGAAACGAGCGAGTACACCCAGCCAAATTACCAATACACCCTGATGGACTGCCAGAATGAGTGCATCCAGCGTACCATAATGGAGACCTGCAGCTGCGTGGACAACCGTATC
AACATGGTAGAGAACCCGCGCCATCTTCCGTTTTGTTTTAAAGTGCCCAGGGTGTCGGAAAAGTGTAGGAACAATACAATGCGTGCGCTGGTAACGAAGACTCAATCGCAGGACGATGAGAAATGTTT
CAAGCCGATCAGGGCGTTCAGTCGGCGCATGGAGTGCCGCAAAGAGGTCTACGAGAACATGACCATCCGCGACCCCGATTCCATGAACAAATGCGAGTGTTACCCGCCCTGCAACGACATTATATACG
ACTCGTCCTATAGTCTGTCAATGTTGCCCGAGCGCAGTCCAGCGCACTCACAATTTTACGCCAGTGTTGTTCGCTTTGTTGAGGATCTTCCAGAACCTCGTCGAATGCTGTTGGAGAACTCTCTGGGC
AAGGACTACAAAGATATTCTCATCAACCGTACAATCCGACTGAATGTGCACATTTCGGACAGCAACATAATCAAAACCACTGAATCCCCTGACTACGAGGCCATTCGCCTGATCTCTGACATAGGCGG
TCAGCTAGGATTGTGGATCGGAATTTCAGTCATGACTATCTTCGAAGTGCTGCAGCTGGTTGCTGACATTTTCCGGTTTCTCACCGGAAAAAGCCGGAACGTTGGTGAACTCCGGAATCGAGCCATTG
ACGATAACTTCCGTAACCATCGCTATGGCAACGGCGACCAAGGATACATACGTCCTTCACATTTGGATCTTGAAAATTGTGAAATCGACAAGCTGACTACAGTGGGATCC
>Capitella_WaNaC_Capca1_212912
GTCGACACCATGGGACTACCTGGATTGCTGGACGCGTTTGCAGGTGGCACCACAATGCATGGTGTCCCCAAAGCCATCAAAGCCCGATCATCTTTTGGACGAGCATTCTGGTCGGTTGTGTGCTTAGC
GGCTGCTTCCATTTTCTGCTTTCAGTTTGCTCAGCTTTTGCGGAAGTACTACTCATACCCAAAGAAGGTCAGCATAGAAATCTTACCTTTGGCCGTTCCATTTCCTGCTATCTCTGTGTGCAATATGC
GTAATTTGGACATCATAGTGTTGAACAAACTCAATGATATCTTCAAGAGAGAGACAAATAATCTCACTGCACAGAATCTCACCGACAACCAGTTCATCAATGAGTACCTCAAAACTGCGTCCAAGTAT
CGCCCAATGGTCCTTCTTCCGGATGTTGACCGTGAGATGTTCCAAACAGTACTCACCAGAACTACGTTCGCTACCAATATAAATCGGGATATAATAGCTACCGGTGGAGTTCCTTTCAAAGAGTTCAT
TGTGACTTGCCGTTTTGGTGGCCATCCGTGTAACCGTACACAAGATTTTAAACACCTGTTTGATCCGTATTACTACAACTGTTATACCTACGAAGCACCTACAACCAGAGTGCCCGAGAGTAGCCTTG
CAGAAGGCCTCGAGAATGGATGGTCAACGACGGTGCTAACCGGGAGTGGAATGCTTGATCGAAATGAAAACATTCGTATTATTCCAGGAACTCATGAGTACTTGAGTCCAATGTCCAGCAGCGAAGGA
GTTCGAGTGGTCATACACCCTCCAAATACGGAGCCATACCCCCACACTGAAGGATTTGATCTGCCACCAGGATATTCTGTAAGTTTTGGAGTCCGCGCTCGGCAGACAATACGCATTGGGGCTCCGCA
TGGTAACTGTTCGCCTAGCAATCCTTTTGGAGACCGGGGTAAGCAGTACCGTTTAATCTCTTGCCAGAAGATGTGTTTGCAGCGAGCAATTGTTGATAAGTGTGGCTGTCGGGATATCGCTCTGCCTG
GCTACGATCAATACCCAGCGACAGTTTTCTGTAATGACGATTCAGTTTTACCTGATTACTGTCGGGAGAATGCCACGGATGAATGCTTAGACGCGGTGAAAGTAATTTATGCCAGGATTATGTGTGTG
AAAGATACGACAGTTGACACAACGCGGAACACAAGTGCAATTCTGGACTGCGGGTGCCACCCACCGTGTAACGAAATCAGTTATGACGTCACTTACAGCCTTTCGAAATGGCCGGCAGAAAGTTTTGA
TGGTGAAGAGGCTTATATCGATATATTTCACGCCGAGAATTACCCGGCACGTTTCACCGGCCCCGAGGACGCCGAGAAGATTGAGCTTTACGGGAACTATTTCGATCCGTCGAATCGGCGCCGTGCCA
TGCAGGACTTTGCCCGTTTGAATGTTTACATAGCTGATTCCAATGTCCTCAAAACTGAAGAGTCCGAGGATTACGTTCAATCGCAGTTGCTTTCGGATCTGGGAGGACAGCTTGGATTGTGGGTTGGG
ATTTCTGTGATTACGATAGTGGAGATTCTGGAGTTAGTCGTTGATTTGTTTCGCTTCCTGTCCACTCGACATGGACCCTACAGTCGAGGAAGTACGTTCTCCAGAGACCCAGAACCCAAGCATCCCGA
CGATGTGTGCCAAAAATGTGAACACTGTAATGGAATCCGAACCACCTGGTTTGACTCTTTTGTTTCCTTCGGAATGCTTCAGAATGCTTGGGGTGGATCC
>Malacoceros_C29205.1.T1
GTCGACACCATGGAGTATACATACAGTAACAGTGATAGGGTGAACAACTCGCCTGGTGGTCCGACACCGAACAACAGGACCCCACAGCACACGAAGATGACTCTATCCAAGGAGAAACTTCACGAACT
GGAACAAGAGTTCCTCCACACCAAGACCAAGGACATTGTCAACAAGTTCATGACCACGACCACTGCGCATGGCTGGGGACGTGTGGGCTCGGAGAATAACATCTTCATCAAGCTGTTCTGGGTGGCTA
TTACACTGACAGCTTTTGGGGCTAATGTGACGCATGTGTTGACTCTGGTGTTGCAGTACAGGGCATTTCCGTCGGAACAAGTGTCTGATGTCGAGCTGTCCTCCATTGAGTTCCCATCTGTTACGATC
TGTAACATTCAACCCATGTCTGTCAGTACTGGCCTGGAGATGATGCAGGACCCATCCACACAGCTCTATCAATGGGACAACCTTACACAATTCTACTATGACATGGCCAAGAAGGACGGTAACTTATC
CTACGAAGAGGAGTTGAAATACAATCGTCTCAAGCAGCCAATTGGTTACTTTGAAAACATTGGTGATGAGGCCACAAGGGTAGGTCATCAGCCCCGAGATTTCCTCCTTAGTTGTACGTTTGGATATA
GGCAATGCACTTGGGATAACTTCACATTCTTCCAAAGCCCCACATATTATAACTGCTACACGTTTAACGGTAACAAAAGTAGTGAGAATCTAGTGGCAAGAACAACTGGACCACAAGAAGGACTGAGT
CTGATCCTTTATCTTGAAAGTGACAATGGTGATGAACTATACAATGGCACATATCATACCTTGTCAAATGTCGGCAATGCAGCAGGTGCTCGGGTGATAGTTCATCCACCTAATTCCAGACCCAGTCC
AGTTGACCAAGGTTTTGATGTACCTCCTGGCTTTTCTAGCTCTGTCGGTGTCAAAGCTACCCGTTACGAGCGACTCGGTGAACCATATGGCCCATGCGTTGAAGACACCATGCATGGTAGTGATCTCT
TTGTGTATGCCACAGACACATGTGTTACTTTATGTCAGCAGCGCTATGTGATGAATGCATGCGACTGCGTCAGTTCCCTTCTACCCATTCCAGAATATAACGAATCAAATTTAAAATACTGTGGGAAC
TTTGACGCAGCACACCAGGAATATTTCTTCTCAAATCTGTCATGTGAAGCGGCATCATTAGCTGAATTTGTCGCAGAAGATAACGTGCGAGAAAGTTGTGGCTGTCATCCACCATGCGAGGAATATTC
ATACAAAACTGACGTTTCATATTCATATTGGCCACTTGACTTCACCCAACTCAGTTTCTATCAGAGCTATGTATTAAATCATCCAGACGTGGACAATCTCAAAGCGTATCAAAATCTTGAGAACTACA
ATACATCAGAGTTGATCTCGACTGGTCTCATACGAAAAAACTTCTTGAGACTTAACGTGTACCTGAAGGATCTGATCATTCAGGAATACATTCAGAAGAAGTCGTATGAGATCCAGAATCTGATCAGC
GACATGGGTGGGACGTTTGGCCTTTGGATCGGCATGTCTGTGATAACATGGTGCGAAGTGGTTGAGCTGCTGATCCGGTTAACATCAAGGAGTGTTCGAAAACTTGTCCACAGCGGAGAGATGCAGGG
TTCGCCTAAGCATTCACAGCGTGGCTCAGGGGTGAACTCTCCCAATAGTATGGACAACAAGGTGTCGCCACGGATGGACCCTCATACTGCCCATGCGCTCGGACTGCAGGATTGTATTGTGTATAAAG
AGGTGCAAGTGGGATCC
>Malacoceros_FaNaC_C40284.0.T2
GTCGACACCATGCCTATCAAGGACGTTTTGGAGAGTTTTGCCAACCACACGAGCATGCATGGCCTGCCTAAGGTCATCAACGCACGGTCAACACTCACAAGATGCTTTTGGGCGGTGATTTGCGTTGC
TGCGGGGGCAATGTTTTGTATGCAAATGACTGAAGTGCTCACGCGCTACTTCTCATACCCCAAAAAGGTGACAGTAGAAGTGGTGCCCACCCCAGTGCCCTTTCCCGCAATATCAATGTGCAACATGA
GAAACCTCGACTTTGACGTTCTCAACACAATCAATCGTAAATTCATCGAGGATCACAGGCCACTGAACCACATCAACACATCAACCAACCCTTTCATCCGCGAGTACATGAAAACAGTCGCCAAATAC
GGTCCACTGTGGTACGAGTACCAAGAGGAGTACCCGCTGATCTTCCAGGAGGTGTTCTCGAGGACTACATTCTCATCCAACATCCCCGAGGACATCATATCATTAGCAGCAGTCCAGCTTGATGAGTT
TGTCGTCAATTGTCATTTTGGCGGCCATGGTTGTAATCGTACGAGAGACTTTGTTAAATTTTTCGATCAGTATTATTTCAATTGTTTCACCTACACAGCCTATCAGGATAAAAATGAAGATGAGTTTT
CATTATCCGAAGGTATCGAAAACGGTTGGTCGGCAATACTATTTTCAGGAAGCAATATGTTGGATAAAAATGAGGATATTAGAGTGTTACCTGGGTTACATGAGTCGCAAAGCCCCGTAAGTGCTAGT
GAGGGAGTGAGGGTTGTTATACATCCTCCAGATACCCAACCATTTCCATTTACAGAGGGTTATGATGTTCCCCCTGGCTTCTCCGCATCGTTTGGAATAAAACCTCGCATGAATATACGCATCGGGCC
TCCACATGGCAATTGCACTAACAAGAATCCATTCGGCAACAAAAACAAATCTCGTTATCGCGGCATCTCGTGCCAGAAAATGTGCTTGCAAAGCCATGTGATCAAAACCTGCGGTTGCTCCGACTCTT
CCCTCCCAAAGGTGCCGGATACCGACGTGAGGCCGTGTAGGAATAATGATGATTTCCCTGATTCATGTATGTTTAATGCCACAAGTGAATGTCTTAGAATACTATTTCGATTGTATGCACGGATACAG
TGCGCTCGGCAGACGCGTGACTGGATAACTAAAAACACAACATTACTTGTTGATTGCCAATGTTTCCCCCCTTGTGATGAAGTGCTTTATGATGTATCTTACAGCTTGTCTAAGTGGCCCGCATCCGG
ATATGAGGGGGATGCTGCCTATTACGATGTATTTTATATTGAGGGTTTTAAGGAGCGGTTCAAGACTACGCCGAAATACGAAAAGATCAAGACGTACTTTAACGACTCGACGCGAGAGACTACAATGA

AGGATTTTGCGCGACTCAACGTCTACGTAGCAGACAGTAATGTGATAAAGACACAGGAATCTGGAGATTACACAACAAATCAGCTAGTGTCAGATATCGGAGGCCAGCTTGGATTATGGGTAGGTATC
TCCGTGATTACGCTGACTGAGGTGATAGAACTGTTATGCGAGATTATGCGCTATCTATTGTCAGCCAAAATGACGACCCCGAAAAGAGACATCGAGCCGGCCGGGGATCGCATGAGAAATGGGAAATA
TAGACCGGCTGCCATGGAGGGACTTCCCCCTACACAATACGACGGCTACGATAGCGCGGATCTTATGTTGGCGTACTATCCGGACCCGCTGCGTAGGGGATCC
>Malacoceros_C44703.2.T1
GTCGACACCATGACATTAAACGACAGTTTTCTCAGTGCGAAGGTGGCATGTTCTAAGTCCAAAATGCCAGCCCCTAAGGATGTTCACCGTGTGTTCCATGAATTTGCCGACATCACATCTATGCATGG
CGTGCCACGCATCATCAACGCTCGTTCCGTGCCTGCTCGTATCTTCTGGTCTGTTATATGCTTAGCCGCATTCGGAATGTTCATGTGGCAAAGCGCCATCCTCTTGCAGCGCTATTATTCCTATCCAA
AGAAAGTTAACATGGAGATCATCCAACGTCCAGTCCAATTCCCTGCCGTCTCAGTGTGCAACTCTGACCCGCTAGATTTGCTTGTGGCTGATAGGCTAAGTGCATATCTTGATGAAAGGAATACCACA
TACAATATAACAGACGAGGAAGAAGATATACTGGTGGCAATTGAAAACAAGTATGGATCATTCTTTACAGCAACAATGGGCTTCTTCCAAGTTTACTCACAGAACATGCAAGCAAGAGGAGGAGAAAT
GAAAGCTGATAGTGTGATTTTAGAAATAAGCTCTCGCTTAGGGCTAACTGCAAATCTAGGGAGGAATTTATCATCACTAGCAGGAATACAATTAAGAGATTTTATTGTAAATTGTAGATTTATGGATG
ATGAGTGTAATATAACAACTGCATTTATGAGAATATTTGATCCCTATTTCTTTAATTGCTATACCTTTAGACCGGAGACTATACTTCCGAGTCGTGCCACCCGTTTGCAAGGTGTGGAGTATGGTCTT
TCATTGTTGCTATTCACAGGAAGCGCAGGTCAGCTTAAGGTTGGAAAAGAGTTGGAGGGTCTGATCATTCCAGGTATGCAAGAAACAGATATCGCATTAGCCTCTGGGCAAGGGGCAAGAGTTGTAGT
GCATTCCCCCCACACTCTCCCTCACCCTGCGGCAGCTGGCTTTGACGTGCCGCCCGGGTATAGCATAGAAATAGGTGTTAAGGCCAGAGAAAACGTGCGCATTCGACATCCGCACGGAAACTGCTCTG
ATCAAGCAGCTAACAAGTCCAACTTCCGTTACACACTCATTCAGTGCCAGCAGGAGTGCATGCAAAATAGGATAATGCAATCTTGCAACTGTGTGGACAATAGGATACCCGAGCCAGAGGATACATTA
GGCTTACGCTATTGCTTCCAATTGCCGGAGTTACCTGAAACGTGTCTGTTCCAACCGTCCTCTCAGCTCTGCTTAAATATCATCCGAGATTGGACTGACAAGATAGATTGTCGGAAAGAAGTCTACGA
AAACATGACAATAAGGGACCCGGAGGCTATGGATAGTTGTGAGTGCTACCCGCCGTGCAACGATATTGTATATGACACGCTTTATAGCCTGTCAACGCTTCCCGAGAACACAGGCGAGCATTCAACCT
TTTATTCCGAGATAGAAAAGTTCCAAAAACTCGGGTTATCCAGTGCTAAAATCGATCACTTGAAGTCACGGTATGGCACTGACTTTGAGACTAAAATGGCAAGTCATATCAGCCGCTTAAATGTTCAT
ATAGCTGATAGTAATATCATCAAAACTACCGAGGCACCTGATTATGAAGCAATTAGACTTGTAAGTGACATTGGTGGACAACTTGGGCTCTGGATTGGTATATCTGTGATGACACTTTTTGAGGTTTT
GCAGTTGTTTGCGGATGTGTTTCGGCAGCTAACAGCCACAGGTCGTCACATAGCCGAGAGAGTACCAACACGCGAGCCCATCGTGGCCACCAGCCTCAACAACAGCTACGACCCTCACGAGCCCTATG
CAACAGACCCTCGGTTACGCGCAAACATAGAACAAGGCATCCCGACCACGAGACGCTTTCACAATTACAACAACGATCGGATAGTCGGTGGATACGATCTGGATATAAATCAACGCTACGAGGTGGAC
AAGCTCACCACTGTGGGATCC
>Malacoceros_FaNaC_C52466.0.T1
GTCGACACCATGGCGATTCGTGATGTAATGACAAAATTTGCTGAACAGACTACTATGCATGGCGTGCCAAAGGTCATTAATGCCAAATCAAGCATGGGTCGTTTGTTCTGGAGTTTAGTCTGTCTGGC
AGCCGGAGCTATGTTTTGTCTACAAATGTCTGAAGTTTTGCAGAGATATTTCAGTTACCCAAAGAAAGTGACTGTTGAAGTCGTCCCCACTCCAGTGCCGTTTCCAAGTATCAGCATCTGCAACATGC
GAAACCTAGATGTTCACATTCTCAACACTCTCAACCGAATGTTCATAGAAGACGACAGACCATTCAGCAATATCAATAAATCGGAACACGAATTTATCCGAGCTTACATGAAAAAGGTGGCTAAATAT
GCCCCACTCTTTTGGAATTACCAAGATGAGTACCCAGAAGTCTTTCAAGAGATATTTAGTCGAACTACTTTCAGTGCAAACATTGATCCGGAGGTAATCGCATTGGCCGCTGTTCAGCTTGAGGGCTT
TGTTGTCAATTGCCATTACGCTGGACATCGGTGCAATAAGACTCGAGATTTCTATCGGTTCTTTGATCCATACTATTTCAATTGCTTCACTTATAAAGCTCACGAACCGACAGACATCGAGGATAATT
TATCAGAAGGGATAGAAAATGGGTGGTCATCTATTTTACTAAGTGGGAGTGGCATGTTGGATAAGAACGATGAGATTAGAATGTTGCCAGGATTACATGAGTGGCGAAGTGCTGTATCTGCCAGTGAA
GGAGTAAGGGTTGTTATACATCCACCGAGTACAACCCCGTACCCATTCACAGAGGGATATGATGTTCCACCCGGGTTTTCAGCTTCATTTGGCATTCATCCTAGACGTAATATCAGAATAGGACCTCC
ACATGGAAACTGTTCTGATAAGAATCCATTCGGTGATGGAACTGAGCGGTACAGACTGATGGCATGTCAGAAGATGTGCATGCAACATTATATTGTAGAAACATGCGGTTGTGCGGATGTTGGCCTCC
CGAAACTACCCTTACAAGCTAATATCTCATGGTGTAGAGACGATGATAACTTTCCTGATGAGTGTATGTTTACCGCTTCAGAGGAGTGCTTACAATTGCTAATGCAACTGCACAACAGGATTAAATGT
GCTCGATCAATAAAATCTAAAATTACTAAAAACACAACTGCCATGGAAGCATGTAATTGTTTCCCACCTTGTGATGAGGTCAGCTATGACGTGAGTTATTCACTTTCAAAATGGCCGTCAGCAGGATA
TGAAGGGGATGCTGCATACTTTGATGTATTTGGAATTGAAAAGTTCAATGAGAGGTTCAATAAAACTGGCACACAGGGCAAATATGAACTATTTACAAAGTATTTCAATGTCTCAAATCGTGAGGAGT
CAATGAAAGATTTCGCTCGCTTGAATGTTTACATAGCTGACAGTAATGTTGTGAAAACTCAAGAATCTGAGGATTACACAAGAAATCAATTGGTGAGTGATATTGGGGGTCAGCTAGGTCTCTGGGTC
GGCATATCTCTTATCACCCTAGCCGAGGTGCTTGAACTAATTATTGATCTGTTTCGTTTATTCTCCAAACATACCTACCGAAGCGTTCCAGTCATTCGACAAAGTATAAAATATAAAGACAAGCGGAA
CGGTGCAGAAATGAACTATGACACAAGATACAGTCAGTCAAATGGCGGACCTCATGCTCGTTATTTACATCATGGACACTCAATACCAAAGCATCCGCCAGAGTTACCTGATACAAGTTTGGATCC
>Malacoceros_WaNaC_C53795.0.T1
GTCGACACCATGCCGCTCGGTCATATGATGCAGATGTTTGCCGGGGGCACCACAATGCATGGTGTCCCCAAGGCAATTCGCTCAAAGTCGTTAACCGGACGGATTTTCTGGAGTTTTATCTGTTTAGC
AGCCGCCTCGATGTTTTGCTTGCAGTTTGCGCAACTTTTGCAGAAATATTACTCGTACCCTAAGAGAGTAACTATTGATATTGTCCCTTCCCCTGTGCCATTTCCAGCTATTTCGTTATGCAACATGC
GTAACTTGGATATCATCGTCCTCAACAACCTTAATAAGATCTTTAAAGAAGCAGCTGATCCCTTAAATTGGGGGAATTACACATCTGATCCGTTTATCAATGCGTACATGAGCGTGGTTGCTAAATAT
TATCCCATGTTTGTCCGGCCGGATATAAACATGCATGTCTTTCAAACCGTTCTCACACGCACATTGATTGCGACTAATATAGAGCGCGATATGTTAGAGCGGGCAGGGGTGCCATTTAAAGAATTTGT
AGTGACGTGTCGCTTTGGGGGCCATGACTGTAATAGGACCAAAGATTTCCGTCAGTTCTTCGATTCATATTATTACAATTGCTTCACATATATTTCGCCATATACCCCTCAGGATCATGATAGTATTT
TAGCTGAGGGGCTCGAGAACGGATGGTCAACGGTTGTTTTGACTGGAGCAGGAATGTTGGATAAAAATGAGGAGATTCGAATGATCCCAGGTACGCATGATCGAGTGAGTCCGATGTCAAGTAATGAA
GGAGTGCGGGTTGTCATACATCCCCCAGACACTGAGCCCTATCCACATACAGAGGGATTTGACGTTGCCCCCGGTTACTCTGTTAGCTTTGGGGTTAAAGCGCGTGAGAACATCCGCATCGGCCCGCC
GCATGGCAATTGTTCCAATAGCAATCCATTCGGCAACGAGTCTTCCGCCTACAGACTCATATCTTGTCAGAAAAAAATGCTTGCAAAAATATATTGTATCTCAGTGCGGGTGTAAAGAGATCTCAATAC
CCGGGCACGATGCCATGTTCCCTGATTTAGACTACTGTACATCGGATAAAGACATCCCTGATCACTGCCGGGAATCGTCTGATGATGATTGTGTTAATGCACTCTTCAAAGTGTATCACCGCTTCCTC
TGTGTGCGTAACGCATCGAAGCACATGACGACAAATGCATCCGCGACACGAGAATGCGGCTGCTACCCACCTTGCCATGAATCAAACTATGACATAACTTACAGCTTATCTAAATGGCCGGCTGAAAG
TTTTGATGGAGAGGAGGCTTATATCGATATATTCGAAACTGAGAACTATCCCGGTAGGTTTATGGAGCCGGAAGATGCACAGAAATTCGATCTTTATGCCAACTATTTCGATGCCTCAAACAGACGGA
CTGCGATGAAGGATTTCGCTCGTTTAAATGTGTATATAGCGGACTCGAATGTGCTAAAAACTGAGGAGTCTGAAGAGTATACACAGTCTCAGTTATTGAGTGATATAGGGGGACAGTTAGGGTTATGG
GTGGGGATTTCTGTAATTACACTTGCTGAAGTATTAGAATTAATTGTGGATGTTGTAAAGTGGGTGGTGCTGTCACATGGACCATGTTCACAGGGTCGGACATTCTCCAAAGACCCTGAAATCGGTGA
ACAAGAACGGGGACTAAATGGGGAAAGTAAATCAAAATCAAGGAACTCAAAGTCCTCAAGAAACTCAAAACGTTCAAACGGGAAAAACAAGAAGCGGCAAATGAACGGCACTATTCCGCTCACGGCTA
CCACCGTGTATGAAGAAGAGAATGAGTACCAGGAACCGGTCATGGATCC
>Malacoceros_C54557.0.T1
GTCGACACCATGGTTGTGTTGTCCATGATTCGCGATTATGCCAGCAACTCAACAGCTCACGGCCTCCCTAAACTTGCCAAAGCGAGAAGCGCGCGTGGGAAGGTGTTCTGGACATTGATATGTTTTAG
TGCTCTGTGCTTTCTTCTCATACTTGGTACATCACTTCTTAGGAAATACTATGCCTACCCTAAGTCAGTTCGTCTGGAGTTGGCAAATCAGCCTATTCCCTTTCCGGCGGTCTCCCTCTGTAATCTGC
GACCAGTAGATGTCTATACCATATGGAAAGCTATCTTTAATGAATCAGGTGAGAGACAATGGGACATCGGAGAAGACTTTGATTACGACGGACGAGATCCACCCGATGCAGATACCTCCATCTTTGAA
TCACATCTCATACGCTTTGCAAACAACTTCAAGTCCTACCAATCCTACTCAGACATCAAAAGTGAGAAGGACCCTATCAAGCTATGGAAGATGGAACGAGAGTTATTGAGTCGTATGACAATGATTGG
GAATCTAAATCTCACAGCTGCACAACAAGGAGGGGTGCAAAAGGAACAGTTCATTGCCAAATGTGAATATGGGGGTGAACATTGCACAGCCAAAAATTTCTCGCTGTTCCAAGATCCGGCCTATCTGA
ACTGCTATACATTTAATCTATCAAAGAACCTGGATTCTATTTATGAGGGTCCTGATAATGGATTGTCTGTTGTTCTCTTTGTCCCTGGCCCATCAATTATGGGACAAGATATGGATAAAGCGGCTGAT
ATTGCCCTCCATCAGGAATTGAGTCTTGGCGGGGAGGGAGTTCGTGTTGTAATACATGAACAAAACACAGTCCCTTATCCACTCACTGAGGGGCTTGACATCCCCCGGGGTGTATCAGCCAGCGTAGC
TATTAAATTGATAGAAAACGATCGACTCGGCCCGCCTCATGGTAACTGCACTGACAAAAAGACGATTGATGGTTTCATAAAATACAGTTACACAATGGCGTCTTGTAAGAAAACATGTCTTCAGAAAC
TTGTGATGGAGACATGCAATTGTGGTGATGTCTCCCTACCCATTTGGAACACCAACCTTACCTTATGTACGAAATTTGATGAGTTACCTGCTGAATGCCAAGGGAGGGAAAATATAACAAGGAATATT
GCGCATTGTGAGGTCTTATTTGATGAATGGTTCCAACGGGTAGGTTGCACTAAATCTACTAAGGCAAATATATCCAAGAATTTATCTGCTTGGACTGAGTGTGATTGTATGCCTCGTTGTCATGACAC
TGAGTATAGTTTATTCTACAGCCTCTCTGATTGGCCGATGCAAGAACAACAAAGAGATGTTGTCCAGGAACTACTTTATGTCGATCGCTTCATCCACAACTTTCCACCAGAAAAACAAAAGCAGTACT
TTGGCTCCATTGATGTAAGAAATAGTTCCCCAACTTATGATGACTACCGCACATTTGTCGAACAGAAAAACTTGATCCGATTGAACATCTACATTTCTGACACCAGTGTTGTGAAAATAATGGAGACG
GAAGCCTATAGTCTGACTCATTTGGCGAGTGATATTGGGGGACAGTTATCTCTCTGGATCGGGGTGTCTATTATCACTATGGTAGAGATAATTGAGCTTATTTGGGGGATGATCAAAATCTGTGCAGC
GGGGACAAATAAAACAAAGGAAGAGGAGAAAGTGAATGGAACAGACATGGAAAAACAGAAGATGAACTATGGGGATGAAGTTGGATCC
>Malacoceros_C67473.1.T1
GTCGACACCATGGATGACACACGTATACAAAGAGGGGAAGTTAAAGGAACACTTTCTTCCTTTGCTGAAAGTACAACTGTTCATGGTCCGTGCCATATCGCACAAGAGAGCAATAAAATTGGCAAAGC
ATTCTGGGTAATAGTATTTCTTGCTGCACTAACTGGTGTCACTATACACCTTTACCTGACTCTGACCGTGTATTACAGCCATCCAACTCATCAATCTCTTTATATCGCTCAAGAAGCTCCGCAGTTTC
CCCACGTGACATTCTGTAATGAAAATCCCATCTCTCGAGAAAACATGGATGAATATGTTAGTATTAATCCAAACTCGAATACAACATGGTTTAGAGGACAGATTGCACAACTCTCATCAAAATTCCAA
GAAGTTGTGGAGCAAATTCCACACATGGATGATTTATTTGCCTCAGCTGCAAGCTACTTTGAAAATATAGGGCGCCATGAAGCCACAAATATTGGACACAAATTTAAGGATTTTGTGATTGATTGCAC
ATATGCAGGAATACCATGCAGCAAAGAGGATTTTGACTTCTTTTTACATCCAATATATTTTAACTGCTATACATTCACAGGGAAACAAGTTAAAAAGAAAGCACAAGAAACTGGTGGAGGTCCTTTTC
TTGGTCTCTCAGTTGTACTGTTTCTTGAGAAGAACCCATCACAGATTCAACTGTACAATAGATATTCTCCAGTTGGCAATGTTGTTGGAACAGTTGTTCAGTTACATTCTGCAAATGACTTTCCCACC
CCTCACATAGAAGGTGTCCATGTTCCTCCTGGTCATTCCTCTGCCATAGCCATGCACACAGAACAACTCAACAAACTTCAACCTCCATTTGGCAATTGCTCAGCTAGACTCCTGCATACACCAGCAGA
TTACCACGTCAGCTACACATCCTCTCTATGTCTGGCACTTTGCCAGCAGAATCTCATCTACCAAGACTGTGGCTGTTACATGGCTACTCAACCTTTCTATGGTCTTGAAAACCCAACATGTGGTACTT
TCCATCTGAATGCGAGTGACAGTGAGAAACTTAGTGTGTATGCCAAGAACATGAAGTGTCAGAATGAAGCCAGCAGGAATGCCAGACTTCGTAGTAGAGATAGGAATATTTGTGACTGCCCACCAGAG
TGTAATAAGACTATTTACTATAAAAGTTTGTCACAAACAGTGTGGCCAGCTGATAATTATTACCGAAATTTCTTGGCTGAATTTCTTGATGAACGACAAGATAAGGCTGACCTTATAGCATACAGGAA
CTTACATCACTTGATAGATACAACATCTGATGAGGTTGAGCTTACCAAATCCATAAAAGAGCAATTTGCTCGAGTAAATGTCTACTTCTCTGACCTTGAAGTAGTTGTAAGACAGGATGTGAGGTCAT
ATACACTTCCACAGCTGTGGTGTGATATTGGAGGCACTATGGGTTTATGGGCAGGTATGTCTGTCATCACTTGTCTTGAGGTTCTGCAACTTCTTGCTAAATTACTGTCTACATTGTGTAAATCTAAG
AGCAATGATAAAACCGGGATTACTTCCATCAGAAATGCTCATACTGAACTGCCTGTCAAGTCGGATCC
>Macrostomum_Mlig049925.g2
GTCGACACCATGCAAGACCGACTCCGCACGGTGACCGGTGCCGCCCAGGAGGTGGCTGGCATCACGAGAACCCACGGCTTGCTGCACATCTTCCTGTCCAGAGGCACAGCTCGACGGCTCTTCTGGCT
GCTTCTGTTCGTAGCTGCGCTGATCGGCTGCACCGTCCACCTGGCAAAGTTGGTGCAGAAGTTCACCGAGCAGTCCGTTGAGTCGCAGCTGAAGCTGGGGAGCGAACGAGCCCAGTTTCCCGACGTCA
CAATCTGCAACTTCAAGCCCGCTTCTGCTAGCTTAATAAGGTGGAATATTTATCACGACTACCTGAATAAATCCTGGGGCAACTTCGACAACTGGTTCCAAGAGGACACCAAGAAGCGCCCGCAAGAA
TCGGATGAAAGCTACGAAGACCGACTTGAGACGGTGTGCAAGAAGCTCCTCGACGTGATGTGGGTGTCAGGCGACACCAGAGACATCTCCCACCTAGATAAGACCATGCTGCTGAGCTGCCGCTACAA
CAGCGAGCCGTGCAGTCACAGGAACTTCTCGCTTGTCCAGACCAGCCGGTTCTGGAGCTGCTATACGTTCCACCCGGACCCAGGGCAGTCGTCCAGCGGTTCTGGGGGCAGGGGAGCCGAGCTGGACA
TGGTGCTCTTCACCGATGGTTATGCCGAACCGTCTATCCATCATTATGAGTTCCTAAATTCCGAATCTGTGGGAGTCTGCGTCAATCGAATAATTTTGGGTCAGGATGCGACGAACAGGTTGCTGAGA
GGCAGTGATCCTCAGTCAGAAGGACTGCGACTTTTTATTCACGAGGCGCACAGCTACCCGATGAGCCAGACGGAGTTCGTGGACGTGGCCTCAGCCACCAGCACCTCCATCAAACTCAGGCCGATCCA
CAACCGACTCAAATCGACGCCGAGCCGCAGATGCTCCGAGCCTCCAATCGAAACCATCAACTATGTCAGGCACTTCTCCAACGCCTCGCTGAATGTGGTCACCAAGGCCTACTCAAAGTCCGTCTCCG
ATTTCGTAGTCGAGGCACAGCAGAGCATCCTGCACAGCAACTGCGGCTGCTATTCGCACCTGCTCCCGTTCTCGGTGAATACGAGCGACCTGTGCTACTTCGCTCCACCGCAGGAGTGGATCACCCCA
TCGGATGCTGTGCTCAAGAGAATTGACTGCCATGATCACTGGTTTGAGCACGCACAGAGGCAAAGCGATGAGCTGGCGAAGGAGTTCCAGCACCGGATGTGGTGCCAATTCACTCAGCAACGGCCTTG
GCAGCAGAGCCGCAGGTGGCCCCCGTTCAGCGCCATTCAAGAGATCTGGGAGAACCTGATGGTGCCGCAAGTCAGACATGGAGTCGAGTTCCTGCCCGAGGATGGCAAAGAGCATCACATATTCAGGA
ATATGAATCAATCTCGACAATACAGCCAAATTTTACCATTTCTAAACGACGATGACTGCATGAGTTTCTTTTGCAATCACAGCCATAAGGATTACTATTATTGCGATACTTTACTGATTCAGCTGTAT
TTGAAGCAAGCGTCGGGTTGCATCGAGCGGTACACTTTGTCTCGCAGCCTGGCGTCAGTCAGCGTCAGCTTGCAGAGCCCGGCAGCTGACGTTTACAAAGAGAAGGCCGGCTACCACTGGACTGAAGC
TCTGTCCGAGGTCGGAGGAACTCTGGGGCTCTGGCTGGGAATCAGCGTTGTCAGCACCTTCGAGCTGCTCGAGTTTTTCTACATTCTGGCGCAGCGATGTCGCGGCGGAACCACCGGCTCGGAGTCCG
AGGGGTCTCCAGACGGATCC
>Macrostomum_Mlig051885.g1
GTCGACACCATGTACAGTCAGGAATCGAACCGACCAGTGGCGAGCTGGGCTGCCCTGCGCGGCGTGTTCATGCACCTGAACAGGTTCGCTCAGCACACCAGCATGCATGGCGTTGCGCACCTATTTAG
AGCCGACAGCTACGTTAGGCGAGCATTCTGGTTGCTGCTCACCTGTCTCACCAGCATGGCCGGATGCTATCACGTCTACCTGACTGTGCACCAGTTCAGCCTGACTCCAGTTAGCACGCTCATTCTTA

ATGAAGGAGTAGACACCATATTCCCGGACGTGACCATCTGCAATCTGCGCCCGATGCCGCAGTACCTGCTGAACCATGATAGTCCAGCTAGACGGCAGTTGGAGTCCAACATGCTTGACCTGATGGTG
GCTCTTAGGCGAGTCTACGCCAAAGGTGGCGACATATTCGGCACGCAGAAGTACTTTCACTGGTACAATCTGTTGCAGCACTACTGGGCGTCAGTGGACACGGCCCATCTGGGCCACAACCTTAGCTT
GTCGATGGTGGCCTGCTTCTACAAGAAGGCCCCGTGCAGACCAGCGGATTTCAAGCTGGTGCAGAACTCGCACTATTGGAATTGCTGGTCATTCCGACCCAAGGATCGAAGCGTGCAAGGCACCGGGC
CCAATGACGGCCTGAATATAATCCTGTACACCAGCACAATGCCGCTGGATGACTCCAGCCAGCCGGATCACGTGGTAGAATATCCGCCGCCAAAGCATCGTTCGACGTTAAGACAATTGGACACGGTG
TTCGGCAAGGCCGGCATGCAAGTGTCCAGCGGCGTCAGGCTGCTCGTGCACGAACCTGGCACGTATCCGCACGTGTACTGGGAGGGCGCCGACGTTGGCAACGGTTGGAGCGCCGACTTGCGCTTCAA
GATGAAGCGAAACGTCTACGTAAACAGAACGGGACATAACTGTGTTGAAAACTACGGCCATACCGACTATTGGGCGTCGGAGGCCGAGGGAATACAGCGGTTTCGCAAACGAGGCCAGGACTGCGTGG
TCAGGAAGTGGCAGGAGCACCTCATGCGCAAGTGCCACTGTCAATCGACATTCCTTCCGACGCAGGACCGCTCGGCTCTCTGCCATTACCTGCGAGCTGCTGGCGTTTCTGCGCAGAACATCTCCGCA
CCTTTCGGGGCCTACAAGTTTCAACAGCTGAAAAGTTTCAACGATGTTGAAAATCAAGTTCGCACGGAGCTTTATGACCAGTTTGCGTCTGAGTGCGGCACTGTTCAGGCCTGCGAGCAGAGCAGTTA
CACATACTCGATCGCATCTGTTCCTTGGCCTGCGCACACCGACCTGGAGGCCTTCATCCACACGTTTGTTAGCCCGAAGTTCCACCGGGCCCGCTACCAGGGCCAGCAGCTGTTGGATCAAATCGTCA
GATTTCACCGACACCCGGACGGGTCGCGGGACTTCGAGAAAGGCTGGAGCGTCAGCCACAGATTTGTGCGGCAAAACTTTGTAAAACTACAGGTCTTCGCAGAAGAGTCCAAAAGCACGCTTATTCAC
GAATCGCCGTCGTATTCGTTTGTTGAGGTTCTGTCTGAGCTAGGTGGAATCGGCGGCCTCTGGGTTGGCATGTCTCTGGTGACATTCGTTGAGCTGTTCGAGTTCCTGGCCATTCTGGGCATCAAGTG
CACTCAAGTTGCGTCCAGCTACCTTCACTCGCGCTGGACTCGGCGAGCTGGTCAAATGGCTAACGGAGCTTGTCCGGGAAGTGCGAACCACCTGCGAACGCCTGCCTGCACCAGGTCCGCCCAATCAC
CCGACGAAATTACTTCGGGACCGTGCGTCGTCCTCGAGGAAGCATCCAGCGTAGTTGATGAGGACACCGAACTGGTTGGAATATTTTTATGCCCGGAAGCGTCCAGAAATGAGCACCTCCACGACGGA
AATTCATCGAGACTTTCGAGAGGACAGGCAATGCAGCACGTGTCTGTTGTGGGATCC
>Macrostomum_Mlig041003.g1
GTCGACACCATGAAGAACACGCTGCATATGATGCGATGCGGCAACTTGATGGCGGCAGCAATGGCGCCTGGGCTGGGGCCTTGCGCTACGAATATTGGGCTGAACAACCTGCAGCAGCAGCAGCAACA
GCAAGGAGCCCAAGAAGAGCAGCAGGGCAATCCGAACGCCCCCGGCAACGGCGGAGGCAGTCGAATACCCAGACGTCGCGAGGTGATGCTTCGCGGTTACATCAACCACTTCACCAGCGGTACGACGG
CTCACGGGCTGAATCGGGTGAACGCCGAACAGAGCTGGGTTCGCATGGTCTGGGTTGGCATCGTCCTCGCAGCCTCATTTGGCGCCATCGTTCACACTACAAAGCAGGTGAAAACCTACCTGCAATAT
CCCGTCAGCAGTACAAATCGTCAAGAGCCGAACTCGTTCGAGTTTCCCGATTTGACATTCTGCGATCCGCTTAATAAGCGCTTCTTTCACAAGAATGTAATTAGTTACGCGCGCGAGGTAGACGTGTA
CGATGAGGCGCACATGATCTACGGCCTCGTCTACAGCCACTTCCGCACAGATTCGGACTGGCGCAAGGAGCGGCGCGCGATCGAACTCATCTACAACGCTCTGGAGCACTTACTGAACATTACCGACG
TCAACATCCGCCCAATGGATGTTGTACTCTACTGCACATTCGACGGCGAACCGTGCAGTCACACGGATTTTCACGTGTTCTACCACCGGCTGTACACCAACTGCTTCACGTTCAAGCCGAAGCAGCGC
AGCCTCCGGAGCAGCGGCATGGACCGCGGACTCTTCCTACTGCTGTACGTACCGTCAGCCGACGTCAACCTCAAACTGGACAGCTTGGCCAGGCTGGCGTCTGGTCTGGAGAACAACGGCATCCGGTT
TCAGATTCACCAGCGGGACACGATCCCGCACCCGCTGGAGTACGGCATCATGGCTCCGACCGGCACCCTGACAGCGGTTGGACTCGAACAAGTGCGCACCTCCCTGGCTGACACGCCCACGAATCCGT
GCGTTCGCGACACAAAACTTCGGCTATTTGATAATCAGTTCAACTTTATGCGACAGTATGACAGACAGCTGTTCGACTGTATCAAGCACCAGTACATGCGCGAAGTTCGCAAACAGTGCGGCTGCACT
TTAGAGAGTCACGTGATCGCCGAGGAGGGCTTCGATCCTCGCGAAGTCGTGTTCTGTCACGATCTGATCAGCGCGAAGAACGCTACGAGAGTTGGAGCGCTCACCAAGCTCTACAATGAACTGAAGAA
TGTCACGAATACAGCGGGTCTTCACGGCCACATCAAGCTGGGGCTGCTCATGCGTCAGAACGACGCTGAGTTGAACGCTTCGCTGGACAGAATGATCTGCTCCGATAGCGTCTCGGAGAGTGCGGCCC
GGCCGGAGCGCTGCTACGATTCGTGCAACTACAACAAATACGAGTACAGCCTCTCTCAGTGCCCTTGGCCGGAGGACTCGTTCGAGGTGCGCAACTCGCAGATCCAGATGGTGAAGATTGCGCAAATG
CTGGAAGATTATAAACTGCGGTACGGTTCGGCCAATCCCGAGACCGCTCTAGCCGGACTCATGCGGAAGAGCAGTCTCAACATTAGCAGCTGTTTGACCCACGATACGCAGAAAATGCAGGTGCCAGA
CAAGGTTGACTGCCATCGGGTGCAGCTGTTTGTAAGAAGATCTGTTGTGCAGCTCCGCGTCTACCCGGAGACTCTGACTGTGCGTCATACGATTGAGGAGCGCAGCTACGAGCTGGTGAACCTGTGCT
CAGAACTCGGCGGCATCCTGGGCCTGTGGATTGGTTTCAGCATCGTGACTCTATTCGAGTTCGCCGAATTATTCATAATATGCGCCTCATACTACTGGTACCTACTGATCAGCAAGCTGCCCAGCATC
CGCAAGAGCCGGCATCCGCTTCGAATCCCAAAACCGCACCCAGTCGGCCTGCGACGCCAAATTCGCCGAGACATTGAGCTGATCTCCAACTCGGGCCAGGGGAACTCGTCATCGCAGCAGGCGTCCCC
ATCTCTATTGGCAACCGGCAGGCCGGCATGCACAAACAATCGAGCGCCACTGTCATCATCCTCATTTGCCAATCATCATCACCACCATCAGCTCCGGCTGGACGGTAATTCGCAGGAAACTCGGCCGC
TTAGAGATGGCATGGGTGGATCC
>Phoronis_g6004.t1
GTCGACACCATGAAGAGGCAGGGTGAGATGGAGGGTCCCGATGGCTTGCTTACCAACTTCGCTCGGAGCACCTCAGCACATGGCTTAGCCCGCATCCCAGGCACCTCCCAGCCAATGCAACGCGCTGT
ATGGGCACTCTTGGTGACGGCCTTGGCAGCTGCTCTCATATCAGCGCTTGCAATCGTCGTTTCCGTTTACCTTCAGTACCAATACACCGAGTTCGCAAAGAAAGTGGCACGCCCTAACATCAAGTTTC
CTGTCATAACAGCGTGCAACAAAGTTCCTTATGGTCTACTCCAAACAGAGCAAACTGTGAATGAGTTTTTTAGGTCAATAGGGTTGGACGAGCCTACATGTCAAAGTGAAATTTCACCAGGATTGAAT
GATCCCATAATATATCGTAGAATTAGCGACGCCGTTAGACCTAGAGTTCTGTACGAATACAACTCTAAAGTAATGCAAGTCTTAGGACAAAACTTGGACGACTTTATGGTGACGTGCGCCTACCAAGG
ACCACCATGCTTTATTTACAGTAACTTCAGCACGTCTTTATTCAAAGACCCTTACCACTATAACTGTGTAACGCTGAAAGTTCCAGACGAAGTTCAGGTAAAAGTCAACGGTCGTGGAGCAATGGAAC
TCGTGTTTTTCGTGGGGGACAACAAAACGTCGCTTCAAGGTAAACATGTGGACAAACTGGTTACGGATGGAACAGTTGGTATAAAATTAATAATTCATGCAGAGGGAATGCATCCCAAATGGGCTAAA
ACACTAGACGTGGCACCTGGCCATTTGGCCAACATCGCTGTACGACCGCGCGAAATTTACAGATTAAAAGCACCGTATGCCAGTGATTGCATAGAGAATCCACCAGATATATTGGCTGCCAGCACGAA
CACTTCCTATACTTACTCGAAAGAGCTGTGTACATTGAAATGTTTCAACAAACTTGTTGCGGACGCCTGCGACTGCATTCCCGAACCTGTCGTGGTGGATAAATTTGGCTTAGACTACCCGTATTGCG
GAGAGTCTCCCTGCAATTATTCGTATATTTTAGATAAAATTAAATGTCACACGAACCTAACAGAGAGATTACATGAAGGTATGTTGTCTAACCAGTGTCCAGAGTGCAAGTTGCCATGTGAAGAATTG
ACTTATGATACTGATATGCATCTTTCAAAATGGCCTTCGAAGTTTACGTCAAAGACCATCACGAAATATATTTTAAAGAACCACGTGATTCCAAGAAACATTGAGAATGATTCTAGTGCTCTCGATTC
ATATGTTCAACAGAACTTTTTGAAAGTGGTGATCTACTTGGAAGACCTTCTTACAGTTGAGCAAACAGAGCAAGAGTCGATGAACTTAGCCCAGCTTACGTCTAGTGTAGGTGGCGCCATGGGTTTTT
TCTTAGGGATATCAATTGTTACTGTGTTTGAGTTCATAGACCTACTAGTCAAGTCTGTACAGTCATTGTTTAAGAAAAAAAGCGTGGACAAAGTGTTGCCATTCCACAGCAAAGGATCC
>Phoronis_g5063.t1
GTCGACACCATGAACTCAGAGGTAAAAAACTGGAGTGTGAAAGACCGCATGAAAGAGTTCTGCGAGTCTACGTCAGCACACGCACTTGGGCAGACTGTAAGTTCTGGCGAAGTGAAGGCGATATTCTG
GTCCCTTGTGTTCCTCAGTGCGTTAGCAGGATGTGTTTGGAATATCGTACACGTTGTGGAGTCGTACACAAGTTTTGGATTCAGCGTAAAGTCAAAACTTGAAATGGAACCGTCTTCACTGAAATTTC
CAAGTGTAACTATTTGTAACCTGAACCCAATAAGTTTCTCAAGACAAGCTAGATTTGCGGGAGACTTTGAAAGGAGTACAGGTTTAGATGATGGGGATTGCGATAATGAGTTTCACAGAGATGAAACT
TATCGTCCAGTCGTCTTGGATTTGTATAGGTGGTCAGAGTTACCCCAATTTTGGTTTGAATACAACGAAAATGTCACAGAAATGTTCGGCCACTCAAAGAAGGATTTTATTGTGGATTGTGTTTTTCA
AAGAGAAGGTGGATGCGAAAACAACTTCTCTGTTACTAAAGACCCCAATCTGTACAACTGCTACACACTGGAACCGAATAAAAACGAAGACCTTCTTGAAATAGGATTTGCAGCTGGTTTATCGTTAA
CTTTATTTGTGGAAAATAGGCGTATTCGGAATGCATTTACTGGCAATTATGCGTTAGATTCCTACCACACTGGAATCGTGGGTGTCAAGGTTGCCATCCACGTGTCGGGTTCTCATCCAAATCCTAAC
ACTCGGGGCGTGATAGCAGAAGTTGGTAAAAGCACCGACTTTATTCTGCGAACAGTAAATAGGACCGCCTTAGGTCCTCCATATCCGTCGCCCTGTAATCCCAAGAAGACAATTGAATCAAGTCATAG
AAAAGAACTTCAGTATGAAGAAAATTTGTGTTTCGCAAGTTGCTTGCAAAACGCCATCGCAAAAAAATGCGGTTGTGTTTCTATCAACCCTATGGCTGTCCCTATAGCAGACAAATTCGGAATAGATC
TTCCATTCTGCGGAAGTTACCCATGTAATGCGACGAAGGTGACAGAAAATTATGAATGCGTCCACGACGTAATAGGCCGTTTCTTACTTAAAAACGACCCCAATTGCCGTAACTGTACCAAGCCTTGT
AACGAAGTCAGTTTTGAAGTTACAAAGTTTCAGTCAAAATGGCCGTCTGAGCTGCACCAGAGTGAGTTCATTAGGTGGCTCGCAAACAAGGACAATACAGCACTGTATAACCAGCTGGTGAGAAATAT
TTCAGATGACAACATTTCCAGATTTATAGAAAACAATTTTCTTCGAATCAACATATACTTTGGGGATTTTTACGTGCGCAAAGACATTGAGACTTTAGATATGGACTGGTTTGACTTGCTCTCAAGTG
TTGGAGGAGCGTTTGGATTCTGGGTAGGAATTTCTGTTGTAACTGGAGTTGAAGTATTGGAGTTACTTCTTGACTGTATTGTCCTTTTTCTAAACAGGCGAATCAAAAAACGGAACCAAATAGATCTT
CGAAACACTACCGGGGACGACGACGTGGTTACGGGATCC

**Other cDNA constructs.** Sources/databases described in *Experimental procedures*.

>Aplysia_FaNaC_BAE07082.1
ATGTTGGGTAGGGGTGAAAGGATAAAGCCTTACCATTTCAGGGACTCGAGCGCAGATCACATGAAATATACGAGCGTCGCGGCCAAGTCGGGAATGGTTCCTGAGCACCGGTACACGATGGTGAGGAG
CCGGCACCACGGCCGCCACCACCACCACCACAGCTACCAGGAGTACAACACGCAGCGCTCCGCCATCAGCCTGATCGCCGAGCTGGGCTCGGAGAGCAACGCCCACGGCTTGGCCAAGATCGTCACGT
CCCGCGACACCAAGCGGAAGGTCATCTGGGCGCTGATGGTCATCATCGGTTTCACGGCCGCCACGCTGCAGCTTTCACTCCTGGTGCGGAAGTACCTGCAGTTCCAGGTGGTGGAACTGTCCGAGATC
AAAGACAGCATGCCCGTGGAGTACCCGTCCGTGACCATCTGCAACATCGAGCCCATCTCGCTGAGGAAGATTCGGAAGGCGTACAATAAGAACGAGAGTCAGAACTTGAAAGACTGGCTCAACTTCAC
GCAGACGTTTCACTTCAAGGATATGTCTTTCATGAACAGCATCCGCGCGTTCTACGAGAATTTGGGCACGGACGCCAAGAAGATCAGCCATGACCTCCGTGACCTCCTCATCCACTGCCGGTTCAATC
GAGAAGAGTGCACCACGGAGAACTTCACGTCCTCCTTCGACGGGAACTACTTCAACTGCTTCACTTTCAACGGCGGCCAGTTACGGGATCAGCTACAGATGCACGCCACAGGTCCGGAAAACGGGCTC
TCGCTCATTATCTCTATAGAGAAAGATGAACCGCTTCCCGGGACGTATGGAGTATACAATTTCGAGAACAACATCCTACACAGCGCCGGCGTACGTGTCGTGGTGCACGCCCCGGGTTCGATGCCCAG
CCCGGTGGACCACGGCTTCGACATCCCGCCCGGGTACTCATCCTCCGTGGGTCTGAAAGCTCTGCTCCACACGCGCCTTTCTGAGCCGTACGGCAACTGCACCGAGGACTCACTCGAGGGAATCCAGA
CGTACCGCAACACGTTCTTCGCCTGCCTGCAGCTGTGTAAACAAAGGAGGCTCATTAGGGAGTGTAAGTGTAAGTCCTCGGCCCTCCCAGATTTAAGTGTGGAGAACATCACGTTCTGCGGAGTCATT
CCGGACTGGAAGGATATACGGAGAAACGTCACTGGAGAATACAAGATGAACCAGACAATCCCCACCATCTCCTTGGCGTGTGAAGCGCGCGTGCAGAAGCAGCTCAACAACGACCGCTCCTACGAGAC
GGACTGCGGTTGCTACCAGCCTTGTAGCGAGACGTCATACCTCAAGTCGGTCTCCCTCTCATACTGGCCCCTAGAGTTTTATCAGCTCAGCGCGTTAGAGAGATTCTTCAGCCAGAAGCACCCGACGG
ACCAGCAGCACTTCATGAAGATCGCCCAAGACTTTCTGTCCCGCCTGGCGCACCCACAGCAGCAGGCACTGGCCCGCAACAACAGCCACGACAAAGACATCCTCACCACCAGCTACTCCCTCTCCGAG
AAAGAGATGGCGAAAGAGGCCTCGGATCTAATACGCCAGAACCTTCTCAGGCTCAATATATACCTAGAGGACCTGAGCGTGGTGGAGTACCGCCAGCTCCCCGCCTACGGGCTGGCCGACCTGTTCGC
GGACATCGGCGGCACGCTGGGCCTGTGGATGGGCATCTCCGTGCTCACCATCATGGAGCTCATGGAGCTCATCATCCGCCTCTTCGCGCTCATCTTCAACGCCGAGCGGGAGGTGCCCAAAGCGCCCA
TGCACAACAGCAACAACGGCGGAAGCGGCGGCGGCGACGGTAGCGGTGGACAGCACAACTTCGCCAACGGGGACGTGGAGCATGAGCGGGACACGCACTTCCCCGACCTCGGCTCCAGCGATTTCGAT
TTTCGCCGCGGCGGCGGGATAGGCGCGGAGTCGCCCGTATG
>Rat_ASIC1a_NP_077068.1
ATGGAATTGAAGACCGAGGAGGAGGAGGTGGGTGGTGTCCAGCCGGTGAGCATCCAGGCTTTCGCCAGCAGCTCCACGCTGCATGGTCTTGCCCACATCTTCTCCTATGAGCGGCTGTCTCTGAAGCG
GGCACTGTGGGCCCTgtGCTTCCTGGGTTCGCTGGCCGTCCTGCTGTGTGTGTGCACTGAGCGTGTGCAGTACTACTTCTGCTATCACCACGTCACCAAGCTTGACGAAGTGGCTGCCTCCCAGCTCA
CCTTCCCTGCTGTCACACTGTGCAATCTCAATGAGTTCCGCTTTAGCCAAGTCTCCAAGAATGACCTGTACCATGCTGGGGAGCTGCTGGCCCTGCTCAACAACAGGTATGAGATCCCGGACACACAG
ATGGCTGATGAAAAGCAGCTAGAGATATTGCAGGACAAGGCCAACTTCCGGAGCTTCAAGCCCAAGCCCTTCAACATGCGTGAATTCTACGACAGAGCGGGGCACGATATTCGAGACATGCTGCTCTC
GTGCCACTTCCGTGGGGAGGCCTGCAGCGCTGAAGATTTCAAAGTGGTCTTCACTCGGTATGGGAAGTGTTACACATTCAACTCGGGCCAAGATGGGCGGCCACGGCTGAAGACCATGAAAGGTGGGA
CTGGCAATGGCCTGGAGATCATGCTGGACATTCAGCAAGATGAATATTTGCCTGTGTGGgGAGAGACCGACGAGACATCCTTCGAAGCAGGCATCAAAGTGCAGATCCACAGTCAGGATGAACCCCCT
TTCATCGACCAGCTGGGCTTTGGTGTGGCTCCAGGTTTCCAGACGTTTGTGTCTTGCCAGGAGCAGAGGCTCATCTACCTGCCCTCACCCTGgGGCACCTGCAATGCTGTTACCATGGACTCGGATTT
CTTCGACTCCTACAGCATCACTGCCTGCCGGATTGATTGCGAGACGCGtTACCTGgtGGAGAACTGCAACTGCCGTATGGTGCACATGCCAGGGgaCGCCCCATACTGCACTCCAGAGCAGTACAAGG
AGTGTGCAGATCCTGCCCTGGACTTCCTAGTGGAGAAAGACCAGGAATACTGCGTGTGTGAGATGCCTTGCAACCTGACCCGCTACGGCAAGGAGCTGTCCATGGTCAAGATCCCAAGCAAAGCCTCC
GCCAAGTACCTGGCCAAGAAGTTCAACAAATCGGAGCAGTACATAGGGGAGAACATTCTGGTGCTGGACATTTTCTTTGAAGTCCTCAACTATGAGACCATCGAGCAGAAAAAGGCCTATGAGATCGC
AGGGCTGTTGGGTGACATCGGGGGCCAGATGGGGTTGTTCATCGGTGCCAGCATCCTCACCGTGCTGGAACTCTTTGACTATGCCTACGAGGTCATTAAGCACAGGCTGTGCAGACGTGGAAAGTGCC
AGAAGGAGGCTAAGAGGAGCAGCGCAGACAAGGGCGTGGCGCTCAGCCTGGATGACGTCAAAAGACACAATCCCTGCGAGAGCCTCCGAGGACATCCTGCCGGGATGACGTACGCTGCCAACATCCTA
CCTCACCATCCCGCTCGAGGCACGTTTGAGGACTTTACCTGCTAA
